# Genetic analysis of global faba bean germplasm maps agronomic traits and identifies strong selection signatures for geographical origin

**DOI:** 10.1101/2022.07.18.500421

**Authors:** Cathrine Kiel Skovbjerg, Deepti Angra, Tom Robertson-Shersby-Harvie, Jonathan Kreplak, Wolfgang Ecke, Alex Windhorst, Linda Kærgaard Nielsen, Andrea Schiemann, Jens Knudsen, Natalia Gutierrez, Vasiliki Tagkouli, Lavinia Ioana Fechete, Luc Janss, Jens Stougaard, Ahmed Warsame, Sheila Alves, Hamid Khazaei, Wolfgang Link, Ana Maria Torres, Donal Martin O’Sullivan, Stig Uggerhøj Andersen

**Author notes:** Authors for correspondence, Cathrine Kiel Skovbjerg (, ORCID: 0000-0001-9780-2920) and Stig Uggerhøj Andersen (, ORCID: 0000-0002-1096-1468), Department of Molecular Biology and Genetics, Aarhus University, Denmark.

## Abstract

Faba bean (*Vicia faba* L.) is a high-protein grain legume crop with great potential for further cultivation. However, little is known about the genetics underlying trait diversity. In this study, we use 21,345 high-quality SNP markers to genetically characterise 2,678 faba bean genotypes. We perform genome-wide association studies of key agronomic traits using a Seven-parent-MAGIC population and detect 238 significant marker-trait associations linked to 12 traits of agronomic importance, with 65 of these being stable across multiple environments. Using a non-redundant diversity panel of 685 accessions from 52 countries, we identify 3 subpopulations differentiated by geographical origin and 33 genomic regions subject to strong diversifying selection between subpopulations. We find that SNP markers associated with the differentiation of northern and southern accessions were able to explain a significant proportion of agronomic trait variance in the Seven-parent-MAGIC population, suggesting that some of these traits have played an important role in breeding. Altogether, our findings point to genomic regions associated with important agronomic traits and selection in faba bean, which can be used for breeding purposes.

**Key Message:** We identified marker-trait associations for key faba bean agronomic traits and genomic signatures of selection within a global germplasm collection.

## Introduction

Faba bean (*Vicia faba* L.) is an important cool-season grain legume (pulse) crop grown worldwide for its high seed-protein content that is of great interest for the production of animal feed and food for human consumption. In addition to an average crude protein content of 29% of dry weight, faba bean shows a high yield potential and is one of the most efficient legumes at fixing atmospheric nitrogen, so it is able to grow in agricultural systems with no applied inorganic nitrogen fertiliser (Singh et al. 2013; Griffiths and Lawes 1978; Baddeley et al. 2013). This provides major benefits for cropping systems and supports sustainable agricultural practices.

In 2020, the total worldwide production of faba bean was 5.7 million tonnes, which shows an overall increase of approximately 55% since 2000 (FAOSTAT 2022). Despite the multiple advantages of growing faba bean, the global production is still surpassed by other pulses such as common bean (*Phaseolus vulgaris* L.), chickpea (*Cicer arietinum* L.), field pea (*Pisum sativum* L.), cowpea (*Vigna unguiculata* L.), and lentil (*Lens culinaris* Medik.) (Adhikari et al. 2021).

In general, faba bean thrives in the cool and moist conditions found in temperate climates, but it is cultivated in various climate zones from boreal to subtropical and warm temperate areas, where it is grown as a winter crop (Singh et al. 2013; O’Sullivan and Angra 2016). Its history of cultivation has been traced back to the Stone Age, making faba bean one of the earliest domesticated crops (Duc et al. 2010). Little is known about the evolution of the species, as the wild progenitor remains unknown. The Middle East is popularly considered the centre of origin, although other studies point towards Central Asia (Cubero 1974; Ladizinsky 1975). Interestingly, no wild faba bean has been found, and *Vicia faba* is not cross-compatible with other *Vicia* species, meaning that all existing faba bean genetic diversity is maintained in germplasm collections and in local populations kept by farmers (Duc et al. 2010). This situation, combined with the current lack of effective transgenic technologies for faba bean, means that ongoing faba bean breeding programmes rely highly on the exploitation of existing genetic diversity.

For optimal crop improvement, it is crucial to obtain a better understanding of population structure and genetic diversity in the accessible faba bean germplasm. To date, the genetic relationships and diversity of faba bean germplasm have been examined in various studies using different molecular marker systems and germplasm collections (e.g. Torres et al. 1993; Link et al. 1995; Terzopoulos and Bebeli 2008; Oliveira et al. 2016; Kaur et al. 2014a; Sallam et al. 2016a, b; Wang et al. 2012; Mulugeta et al. 2021). Although these studies have found genetic distinctions between germplasm belonging to different geographic origins, the underlying selection signatures remain unstudied and poorly understood. This is mainly due to the large and complex genome of faba bean (approx. 13 Gbp) (Khazaei et al. 2021).

The identification of genomic regions differentiated between subpopulations of faba bean with different geographic origins will be an important factor in addressing the challenges that come with the increasing climatic fluctuations of global warming. Signatures of selection have been identified in multiple important crops such as maize (*Zea mays* L.), rice (*Oryza sativa* L.), alfalfa (*Medicago sativa* L.), and soybean (*Glycine max* (L.) Merr.) by comparing subgroups with different geographical origins (Xu et al. 2022; Bouchet et al. 2013; Xie et al. 2015; Chen et al. 2021; Saleem et al. 2021). This is typically done using statistical methods that rely on differences in allele frequencies between subpopulations (Luu et al. 2017; Chen et al. 2010; Tajima 1989; Foll and Gaggiotti 2008). Meanwhile, genome-wide association studies (GWAS) have consistently proven to be a powerful tool for detecting candidate genes for agronomically important traits (Huang et al. 2010; Sonah et al. 2015). By looking for overlaps of genomic regions under selection and quantitative trait loci (QTLs) identified by GWAS studies, it is possible to study traits under selection. However, traits under selection during breeding are typically strongly correlated with population structure, posing a challenge to GWAS. GWAS models correcting for population structure will cause many false negatives, and ultimately QTLs associated with traits under selection might not be identified. On the contrary, a naïve GWAS model with no population structure adjustment will yield too many false-positive signals, since it is not able to distinguish genetic regions associated with overall population structure from causal genes associated with traits under selection (Zhao et al. 2011). A way to overcome this problem is to combine selection signatures of a diversity panel with GWAS results from independent populations. This is especially straightforward for well-studied crops such as maize and rice, where a large number of functional genes and loci associated with traits have already been identified and published (Xu et al. 2022; Xie et al. 2015). For an orphan crop such as faba bean, however, most QTLs associated with agronomic important traits are still to be identified (Adhikari et al. 2021).

In view of the above, the objectives of the present study were to: (1) analyse the genetic diversity, population structure, and linkage disequilibrium (LD) of a global faba bean panel of 2,678 accessions using high-quality SNP data; (2) use a mapping population to identify markers associated with key agronomic traits; and (3) select a large, non-redundant diversity panel to study the genetic diversity and inter-population selection signatures of faba bean. Understanding the genetic diversity and structure of these accessions lays a foundation for future genome-wide association studies (GWAS) or genomic selection (GS) and will aid in the utilisation of these materials in future faba bean breeding programmes.

## Materials and methods

### Plant materials and panels

The data studied consists of 2,678 faba bean accessions belonging to eight different international panels referred to respectively as: EUCLEG, GWB, Four-way-cross, Seven-parent-MAGIC, NORFAB, ProFaba, RSBP and VICCI. Further details on the individual accessions can be found in **Supplementary File 1**.

The EUCLEG panel contains data on 358 accessions from different countries of Africa (17 accessions), North, Central, and South America (8), Asia (60), and Europe (165). The origins of the remaining 108 accessions are unknown. Europe, with 19 countries, is the most-represented geographical area in the panel, followed by Asia, America, and Africa (16, 4, and 3 countries, respectively). Spain accounts for the highest number of accessions (62). The panel includes 55 breeding and advanced materials, 57 breeder lines, 8 inbred lines (parents of hybrid cultivars), and 237 accessions from different germplasm banks, aiming at gathering a wide range of genetic diversity from diverse geographical origins. The EUCLEG panel was made in collaboration with public institutes such as ICARDA, IFAPA, IFVCNS, INIA, and INRA, Ghent University, the University of Göttingen, and the following genebanks: ESP004, ESP046, FRA043, SWE054, and SYR002. Private sector contributors included five seed companies (Agrovegetal, NordGen, NPZ, Batlle, and Fitó). Prior to the genotyping analysis, all the Spanish lines had been selfed for at least four generations. The remaining accessions were purified for two generations by single seed descent (SSD) in insect-proof cages. Afterwards, tissue samples were harvested from a single plant before the DNA was extracted.

The NORFAB panel contains data on 195 accessions with large trait diversity and wide geographical origins. Most accessions have a European origin (85), but other continents are represented as well: Asia (43), Africa (9), North America (7), South America (6), and Australia (1). The geographical origin of the remaining 44 accessions is unknown. The most represented countries are Denmark and Germany, which each contribute 16 accessions. When the panel was established, there was a focus on including accessions with resistance to important pathogens of North European conditions. The material was obtained from the University of Reading, the University of Göttingen, the University of Helsinki, Nordic Seed, seed companies, and genebanks. Materials received from universities and Nordic Seed were inbred for three or more generations and thus considered sufficiently homogenous for direct entry in the core collection. Plant materials from the genebanks and seed companies were inbred by growing 10 seeds of each accession in a nethouse. The offspring of a representative plant within each accession was further inbred for two generations in insect-proof greenhouses or nethouses by SSD.

The ProFaba panel contains data on 234 nonredundant accessions. This panel of global diversity is based on joint donations from breeders (Agrovegetal, Sejet Plant Breeding, NPZ Lembke KG, and GSP) and from academic and research institutions (the University of Helsinki, the University of Göttingen, IFAPA, INRA, the University of Reading, and Teagasc). Most accessions were highly homozygous inbred lines such as Hedin/2, Hiverna/2, or ILB 938/2; a few were partly inbred lines, genetically narrow cultivars, or old varieties such as Alameda or Irena. The initial collection of entries was guided by available knowledge about the global faba bean germplasm. It attempted to represent minor, equina, and major types, European spring and winter types, and also to include germplasm from the Middle and Far East, North Africa, Australia, and Latin America, but it was in practice determined by availability of controlled-inbred seed. The 234 inbreeds included here were retained from a longer initial list of 259 genotyped lines, with redundant and/or heterozygous lines being excluded from the analysis.

The GWB panel contains data on 268 winter faba bean accessions. These are inbred lines (F>9), derived via SSD from the so-called Göttingen Winter Bean Population (GWBP). That population stems from mixing 11 inbred ancestors, which for their part were derived from 11 winter bean cultivars and academic accessions. These 11 ancestral sources were chosen to represent true winter beans (Bond and Crofton 1999) and yet include diverse materials. The sources comprise three German cultivars (“Webo”, “Wibo”, and “Hiverna”, bred by H. Littmann), three accessions from H. Herzog (“79/79”, “977/88/S1/8”, and “979/S1/1”), three UK cultivars (Banner, Bourdon, and Bulldog), and two French types (the landrace “Côte d’Or” and the cultivar “Arrissot”). The 11 ancestral inbred lines were mixed and, according to their partial outcrossing, allowed to recombine across several generations (Gasim et al. 2004), and thus, from 1992 to 1999, the GWB was created. In 1999, the single seed descent procedure started. In that way, the lines that serve here as GWB panel were bred from the GWBP. The current GWB panel is maintained via strict selfing. The panel does not have the formal structure of a MAGIC population because the GWB panel was not created via a crossing scheme but by repeated random partial allogamy (approx. 40–50% cross-fertilisation); and because some natural selection for hardiness was allowed to occur between the creation of the GWBP and the initiation of the single seed procedure (Link and Arbaoui 2006).

The Virtual Irish Centre for Crop Improvement (VICCI) population is a complex outcrossing population consisting of 563 accessions. The population was created between 2016 and 2019 by allowing F_1_ hybrids from many combinations of 22 founding lines to outcross in the presence of captive bumblebee colonies for 6 generations, with recurrent selection for single plant seed yield performed in alternate (summer) generations. The 210 founding F_1_ hybrid individuals and 353 individuals from the first, second, and third generations of selection together represent a longitudinal sampling of this recurrent selection experiment and will be collectively referred to as the VICCI panel (Tagkouli 2021). The key difference between this panel and the other materials presented here is the absence of any opportunity for sustained inbreeding.

The Reading Spring Bean Panel (RSBP) panel contains data on 160 accessions and was developed by deriving inbred lines from selected progeny of the VICCI population. After the first round of selection within the VICCI recurrent selection scheme, up to four progeny seeds of the 54 highest-yielding individual VICCI plants were put through SSD for three generations from 2017–2019. From this process were derived 160 genotyped S_3_ lines, which constitute the RSBP (Warsame 2021).

The Seven-parent-MAGIC panel contains data on 255 multi-parent advanced generation intercross (MAGIC) accessions. It was developed from seven parent lines from the inbred NORFAB panel: Mélodie, Diana, Albus, Hedin/2, Icarus, ILB 938/2-2, and CGN07715. Mélodie is a French variety with low vicine-convicine content and a colourless hilum; Diana is a German variety with auto sterility; Albus is a Polish variety that is low tannin (*zt1*) and has a colourless hilum; Hedin/2 is a German variety with auto sterility and small seeds; Icarus was bred in Australia and, together with ILB 938/2-2, shows a large degree of branching and resistance to *Botrytis fabae*; and the CGN07715 is a closed flower mutation that produces tall plants which are late to mature. The Seven-parent-MAGIC panel was generated by a half-diallel crossing between all seven parents. Whenever relevant, the female parent was chosen as the genotype, carrying recessive phenotypic characters such as tannin free, colourless hilum, and closed flowers. Subsequently, all F1 individuals that did not share any parents were crossed in a nethouse. This produced 102 out of 105 possible crosses, which were all inbred in insect-proof greenhouses or nethouses for three generations by SSD.

The Four-way-cross panel contains data on 645 recombinant inbred lines (RILs) at F5. It was developed from four founders: (Disco/2 × ILB 938/2) × (IG 114476 × IG 132238) (Khazaei et al. 2018a). Disco/2 is bred at INRA (France) as both a low tannin (*zt2*) and low vicine-convicine (*vc^−^*) source. ILB 938/2 (IG 12132) is from the Andean region of Colombia and Ecuador and has resistance to multiple biotic and abiotic stress (Khazaei et al. 2018b). IG 114476 is a *Paucijuga*-type faba bean with very small seeds from Bangladesh, and IG 132238 is an early maturing faba bean genotype from China.

### DNA extraction and SNP genotyping

Genomic DNA was extracted from fresh leaf tissue using a DNeasy Plant Mini Kit (QIAGEN Ltd, UK) for the EUCLEG panel, a NucleoSpin Plant II kit (Macherey-Nagel) for the Seven-parent-MAGIC and NORFAB panels, and a DNeasy 96 Plant Kit (QIAGEN Ltd, UK) for the remaining panels. DNA quality was assessed on agarose gel electrophoresis, while concentration was assessed using a Quant-iT PicoGreen dsDNA Assay Kit (ThermoFisher Scientific, UK) following the manufacturer’s guidelines.

SNP calling was performed using the Vfaba_v2 Axiom SNP array (Khazaei et al. 2021 O’Sullivan et al. 2019). The flanking sequences of 35,363 SNPs were aligned to the *Vicia faba* reference sequence (https://projects.au.dk/fabagenome/genomics-data) using the blastn application of the NCBI BLAST+ suite of programs (v2.12.0+) with an e-value of 1e-8 as the significance threshold. From the resulting alignments for 34,854 SNPs the best alignments per sequence were selected based on maximal bit-score. Sequences with multiple alignments at their best bit-score were removed, leaving 31,186 SNPs that had produced unique best alignments in the BLAST analysis. A further 23 of these had to be removed because they did not include the SNP position in the alignment. For the remaining 31,163 SNPs, the SNP position in the reference sequence was determined based on the “Blast trace-back operations” (BTOP) string of the alignments, counting upwards from subject start to the SNP position in the query sequence when the alignment was on the plus strand and downwards when it was on the minus strand. The significance threshold for the blast analysis had been determined by aligning the first 1,000 flanking sequences to the reference genome without a threshold, selecting the best alignments per sequence from the results by maximal bit-score and taking the highest e-value among the best hits rounded to the next higher full figure as significance threshold. Functional annotation was done using eggNOG-mapper v. 2.7.2 with the eggNOG eukaryotic database (Buchfink et al. 2015; Huerta-Cepas et al. 2017, 2019).

Finally, the set of mapped markers were filtered to include only SNPs categorised as “PolyHighresolution markers” (21,423), and markers that mapped to a non-unique position in the genome (78) were removed as well. This gave a set of 21,345 high-quality quality markers with positions for downstream analyses. Note that chromosome 1 was provided split into two parts (Chr1S and Chr1L) at position 1,574,527,093 by the faba bean genome consortium to facilitate data analysis.

Using the 8,423 markers for which both a genetic and physical position (Supplementary File 2, https://projects.au.dk/fabagenome/genomics-data) were available, we modelled, in the R package *cobs*, genetic position as a smooth, strongly monotonic function of physical position, and then, using this function, we estimated genetic positions for all SNPs (Ng and Maechler 2007). These genetic positions were used for imputation of missing genotypes using Beagle v. 5.2 with windows of 60 cM and 3 cM steps (Browning et al. 2018). All markers showed missingness <5% before imputation.

### Redundancy filtering

Genetic identities (*GI*) were calculated as the fraction of shared alleles using SNP data and the following equation (**1**):

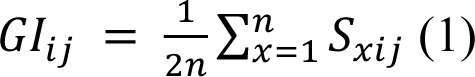

where *GI_ij_* is the genetic identity between the *i*th and *j*th sample, *n* is the number of markers where none of the two samples show missingness, *S_xij_* is the number of shared alleles between sample *i* and *j* at marker *x* and therefore takes values of 0, 1, and 2.

The redundancy filter was applied within individual panels, where 12 accessions were genotyped in duplicates, and to the diversity panel, where the combination of multiple panels resulted in redundancy. When two samples showed *GI ≥* 94%, the sample with the largest proportion of genetic missingness or the least information in terms of geographic origin (diversity panel) was removed. The threshold was set so that we excluded most duplicates of accessions and avoided discarding too many genetically close lines.

### Population structure and phylogeny

To infer the population structure, the software ADMIXTURE was run with K ranging from 2 to 20 (Alexander et al. 2009). A ten-fold cross validation (CV) scheme was repeated 10 times for each value of K. After plotting the average CV error as a function of K we found that the local minimum was reached at K = 15, but that the relative reduction of the CV error when going from K to K+1 was significantly smaller (less than 1%) after K = 4 (**Supplementary Figure 5**). With this in mind and for interpretation reasons, we considered the best value of K to be between 2 and 4. The optimal number of K was then chosen as the value where genetic subpopulations reflected geographic subpopulations to the highest degree. The admixture proportions were graphically displayed using R.

To further confirm the subpopulations identified by ADMIXTURE, we performed principal component analysis (PCA). All PCAs in this study were made by using PLINK v. 1.9 with a minor allele frequency (MAF) filter at 0.01 and by setting the number of principal components to the number of samples (Purcell et al. 2007). The resulting eigenvectors were plotted in R using ggplot2 (Wickham 2016). Accessions were assigned to a subpopulation if they showed ancestry proportions ≥0.50.

### Genetic variation and diversity

The site-frequency spectrums were based on polymorphic SNPs. The alternative allele counts and resulting plots were made using a custom R-script. Nucleotide diversity was calculated by applying “--site-pi” in VCFtools v. 0.1.16 (Danecek et al. 2011) using all 21,345 markers. The heterozygosity of individual panels was calculated in R using the inbreedR package (Stoffel et al. 2016). SNP densities were calculated chromosome wise using ‘--SNPdensity’ with a distance of 1M base pairs (bp) in VCFtools v. 0.1.16 (Danecek et al. 2011).

Population differentiations were investigated by calculating fixation indices (*F_ST_*) between pairs of subpopulations as identified by ADMIXTURE. For this purpose, the ‘--weir-fst-pop’ in VCFtools v. 0.1.16 was used (Danecek et al. 2011). Additionally, analysis of molecular variance (AMOVA) was performed using the adegenet package in R (Jombart 2008).

### Identification of SNPs under selection

To detect SNPs showing signatures of selection, we employed three methods of outlier detection that differ in their statistical approaches. All aim to identify extreme differences in allele frequency between populations.

Our first approach used the software package Ohana with the number of ancestry components set to 3 (Cheng et al. 2022). Ohana works in a two-step manner. First, population structure and the allele frequencies of the ancestral admixture components are co-estimated through an unsupervised learning process. Second, maximum likelihood models are used to identify signals of positive selection on ancestry components. This is done by looking for SNPs that deviate strongly from the globally estimated covariance structure using a likelihood ratio (LR) test. Consequently, SNPs associated with positive selection will show high LR values.

Our second approach applied the R-package pcadapt with K = 3 (Luu et al. 2017). Pcadapt uses a PCA-based approach to simultaneously infer population structure and identify loci excessively related to this structure.

Our third approach used the software BayeScan v. 2.1 with default settings (Foll and Gaggiotti 2008). The software builds on hierarchical Bayesian models and decomposes locus-population *F_ST_* coefficients into a population-specific component and a locus-specific component. It then calculates the posterior probability of a locus-specific component being different from 0. In contrast to Ohana and pcadapt, BayeScan requires grouping into populations. For this purpose, we used the population memberships assigned by ADMIXTURE.

For candidate markers under selection, we chose to focus on markers found by at least two of the methods, thus reducing the risk of false positives.

### Linkage disequilibrium

Linkage Disequilibrium (LD) was estimated individually for each panel using PLINK v. 1.9 to compute the squared correlation coefficients (*R^2^*) chromosome-wise for each pairwise combination of markers (Purcell et al. 2007). Before LD calculations a MAF filter was applied at 5% in individual panels and 1% in the diversity panel. For each panel, the resulting LD data was merged across chromosomes, subsequently sorted according to SNP distance, and binned into groups of 1000 data points. For each bin, the average *R^2^* was plotted against the average distance (bp) and a smooth curve was fitted using the *loess* function with a 10% smoothing span in R. There were 5000 bins plotted for the populations Seven-parent-MAGIC + Four-way-cross where the LD decayed slowly, whereas 1000 bins were plotted for the remaining populations. The LD decay was estimated per panel as the point where the fitted curve reached half of its maximum value.

### Phenotyping and field trials of Seven-parent-MAGIC lines

The Seven-parent-MAGIC lines were grown in field trials at two locations in Denmark during 2020 and 2021.

The first location was at Sejet Plant Breeding, Sejet (55.82°N, 9.94°E) and the second at Nordic Seed, Dyngby (55.96°N, 10.25°E). The setup of field trials was randomised with three replicate blocks. Plants were grown in plots made up of 6 rows with 14–15 seeds sown per row. Each plot contained two entries and therefore consisted of two inbred lines, each contributing three rows. To minimise neighbour effects, every other plot between the 6-rowed plots consisted of commercial cultivars; Kontu at both field trials during 2020 (Sej20 + Dyn20) and Taifun and Daisy at both field trials during 2021 (Sej21 + Dyn21). For the field trials in Sejet, the sowing dates were 9 April 2020 and 19 April 2021, and the harvest dates were 10–12 September 2020 and 31 August to 01 September 2021.

For field trials in Dyngby, the sowing dates were 2 April 2020 and 8 April 2021, and the harvest dates were 24 August 2020 and 21 August 2021.

Trials were rain-fed and treated with herbicides and insecticides. Furthermore, the field trials in Dyngby had fertiliser (NPK 0-8-23) applied. More details on the treatments can be seen in **Supplementary Table 1**.

To minimise border effects, all traits were scored in the middle row of the three rows per inbred line. Plants were phenotyped for the following 17 traits: disease susceptibility to chocolate spot (caused by *Botrytis fabae*), rust (caused by *Uromyces viciae-fabae*), and downy mildew (caused by *Peronospora viciae*); herbicide damage; branching; plant height; number of ovules per plant; sterile tillers per plant; lodging; maturation date; earliness, end, and duration of flowering; thousand grain weight (TGW); and seed area, length, and width. The description of each trait and score appears in **Supplementary File 3**.

### Statistical models and genome-wide association studies

Prior to GWAS, the phenotype scores were filtered for extreme outliers and data that came from lines with signs of mixed identity. This left between 188 and 234 Seven-parent-MAGIC accessions for GWAS.

Phenotypic data analyses were performed for each trait in individual field trials and for all environments (envs) combined, using the lme4 package in R (Bates et al. 2014). For analysis of variance (ANOVA) and to get adjusted genotype means for GWAS inputs, we fitted the following mixed model to all traits (**Eq. 2**):

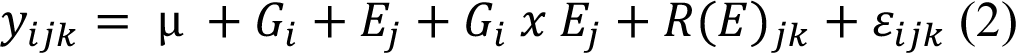

where *y_ijk_* denotes the phenotypic value of the *i*th inbred line in the *j*th environment (year x location combination) in the *k*th replication, *µ* is the overall trait mean, *G_i_* is the genetic effect of the *i*th line, *E_j_* is the environmental effect of the *j*th environment, *G_i_ x E_j_* denotes the genotype environment interaction of the *i*th line in the *j*th environment, *R(E)_jk_* is the effect of the *k*th replicate within the *j*th environment, and ∈_ijk_ is the residual error. All effects except the overall mean were treated as random. If a trait was scored on two separate dates within a field trial, each date was modelled as a separate environment. All random effects in the model were tested one at a time for statistical significance by using the ‘ANOVA’ function in R to compare the log-likelihood of a model with and without the random effect. When testing the significance of main effects, the interaction effects were excluded from the full model before the main effect was dropped. If the removal of an effect was associated with a *p*-value > 0.05, inclusion of the effect was not considered to improve the model. To extract best linear unbiased estimators (BLUEs) for each trait, statistically insignificant terms were excluded from equation 2, which was then refitted with genotypes as a fixed effect. For the trait x environment combinations where *G_i_* and ∈_ijk_ were the only significant effects, the average phenotype value of each genotype was used for GWAS.

Broad sense heritabilities were calculated on a line mean basis from the estimated variances of equation 2 (**Eq. 3**):

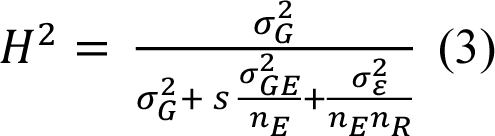

where σ^2^_G_, σ^2^_GE_, and σ^2^ are the estimated variances of the genetic effects, genotype x environment interactions and residual effects, respectively; *n_E_* and *n_R_* are the number of environments and replications, respectively; and *s* is a constant taking the value 0 if only one environment is included in equation 3 and otherwise taking the value 1. It should be noted that when *s* = 0, *H^2^* is strictly speaking a measure of line repeatability and not line heritability.

GWAS was performed on the generated BLUEs using the Fixed and random model Circulating Probability Unication (FarmCPU) method integrated in the GAPIT v. 3 library in R with a MAF filter of 5% (Liu et al. 2016; Wang and Zhang 2021). To avoid signals originating from population stratification, the first three PCs were included as covariates in the GWAS models. Finally, we verified the absence of confounding effects by checking for inflation of *p*-values by examination of Q-Q plots and calculation of genomic inflation factors (λ). For traits where λ-values were not between 0.9 and 1.1, *p*-values were divided by λ. To avoid the high penalty of Bonferroni correction, which assumes all markers are uncorrelated, we calculated the effective number of independent tests (M_ef_) using the SimpleM method (Gao et al. 2010). The significance threshold was then estimated as 0.05/M_ef_ with M_ef_ being 19,697 for the diversity panel and 4,790 for the Seven-parent-MAGIC panel.

The phenotypic variance explained (PVE) by SNPs were estimated for all traits as proposed by Martinez et al. (2018) (**Eq. 4**):

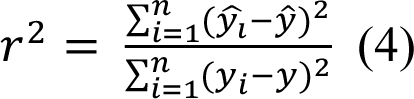

where *y*_i_ is the phenotypic value of the *i*th observation (not corrected for any effects as included in equation 1) and ŷ_i_ is the predicted value of the *i*th observation when phenotypes are fitted as a linear regression of the genotype of the significant SNP(s); *n* is the total number of observations.

## Results

### Genomic distribution of SNPs

We genotyped a large collection of 2,678 accessions from eight different panels (EUCLEG: 358, Four-way-cross: 645, GWB: 268, Seven-parent-MAGIC: 255, NORFAB: 195, ProFaba: 234, RSBP: 160, VICCI: 563). After MAF filtering, the data contained 21,345 high-quality SNPs that all mapped to unique positions in the faba bean genome (www.fabagenome.dk). SNPs were well-distributed across chromosomes, and the average SNP density was 1.9 SNPs/Mbp (**Table 1**). The average distance between two adjacent SNPs was 542.8 kbp.

**Table 1.** SNP distributions and nucleotide diversities

| Chr | Length (bp) | # SNPs | SNP density* | SNP distance** | $\pi$ (nucleotide diversity) | | | | | | Seven-parent-MAGIC | Four-way-cross |
| --- | --- | --- | --- | --- | --- | --- | --- | --- | --- | --- | --- | --- |
|  |  |  |  |  | EUCLEG | NORFAB | ProFaba | GWB | RSBP | VICCI |  |  |
| Chr1S | 1,574,527,093 | 2262 | 1.4 (10) | 705.6<br>(28,576.9) | 0.33 | 0.33 | 0.33 | 0.29 | 0.30 | 0.31 | 0.30 | 0.27 |
| Chr1L | 1,805,244,829 | 3852 | 2.1 (14) | 475.7<br>(31,207.6) | 0.32 | 0.32 | 0.32 | 0.29 | 0.29 | 0.30 | 0.29 | 0.27 |
| Chr2 | 1,716,769,615 | 3633 | 2.1 (19) | 478.4<br>(22,075.1) | 0.32 | 0.32 | 0.32 | 0.29 | 0.28 | 0.29 | 0.28 | 0.25 |
| Chr3 | 1,637,815,978 | 3292 | 2.0 (20) | 509.6<br>(41,013.2) | 0.32 | 0.32 | 0.32 | 0.29 | 0.28 | 0.29 | 0.29 | 0.27 |
| Chr4 | 1,645,877,737 | 3008 | 1.8 (13) | 566.5<br>(60,679.0) | 0.32 | 0.32 | 0.32 | 0.28 | 0.29 | 0.29 | 0.28 | 0.26 |
| Chr5 | 1,365,994,436 | 2611 | 1.9 (19) | 548.8<br>(68,652.8) | 0.32 | 0.33 | 0.33 | 0.28 | 0.29 | 0.30 | 0.29 | 0.26 |
| Chr6 | 1,520,236,431 | 2687 | 1.8 (18) | 597.3<br>(86,906.2) | 0.32 | 0.32 | 0.32 | 0.28 | 0.28 | 0.30 | 0.29 | 0.26 |
| Avg. | 1,609,495,160 | 3049 | 1.9 | 542.8<br>(86,906.2) | 0.32 | 0.32 | 0.32 | 0.28 | 0.29 | 0.30 | 0.29 | 0.26 |

\* Average number of SNPs per 1 Mbp; parentheses include highest number of SNPs.
\*\* Average spacing between neighbouring SNPs in kbp; parentheses include the largest distance.
Abbreviations: Chr, chromosome.

### Characterisation of individual panels

Most inbred panels showed average heterozygosity below 0.06. An exception was the EUCLEG panel, which comprised a group of accessions with higher heterozygosity and therefore averaged 0.14. The outcrossing nature of the VICCI panel was demonstrated by its average heterozygosity being remarkably higher (0.18) than for the remaining panels (**Supplementary Figure 1**).

To compare the genetic diversity captured within each individual panel, we investigated the distribution of minor allele count (MAC) and calculated nucleotide diversities (π). The nucleotide diversity was highest for the broad diversity panels, EUCLEG, ProFaba and NORFAB (0.32), whereas the remaining panels, which were all established from a limited number of founder lines, had lower π-values (0.26–0.30) (**Table 1**). The distribution of MACs was very similar for EUCLEG, NORFAB, and ProFaba, which showed a close-to-uniform distribution with a small overrepresentation of intermediate frequencies. For the remaining panels, we observed an excess of low-frequency variants. The distribution of MACs for the mapping panels, Seven-parent-MAGIC and Four-way-cross, was multimodal and reflected the numbers of founder alleles present. Thus, in addition to the peaks close to zero, the Seven-parent-MAGIC population showed an excess of markers with MAC of ∼75 (frequency of 1/7), ∼150 (frequency of 2/7), and ∼220 (frequency of 3/7), while the Four-way-cross showed additional peaks at ∼325 (frequency of 1/4) and ∼645 (frequency of 2/4) (**Figure 1A**).

**Figure 1.**
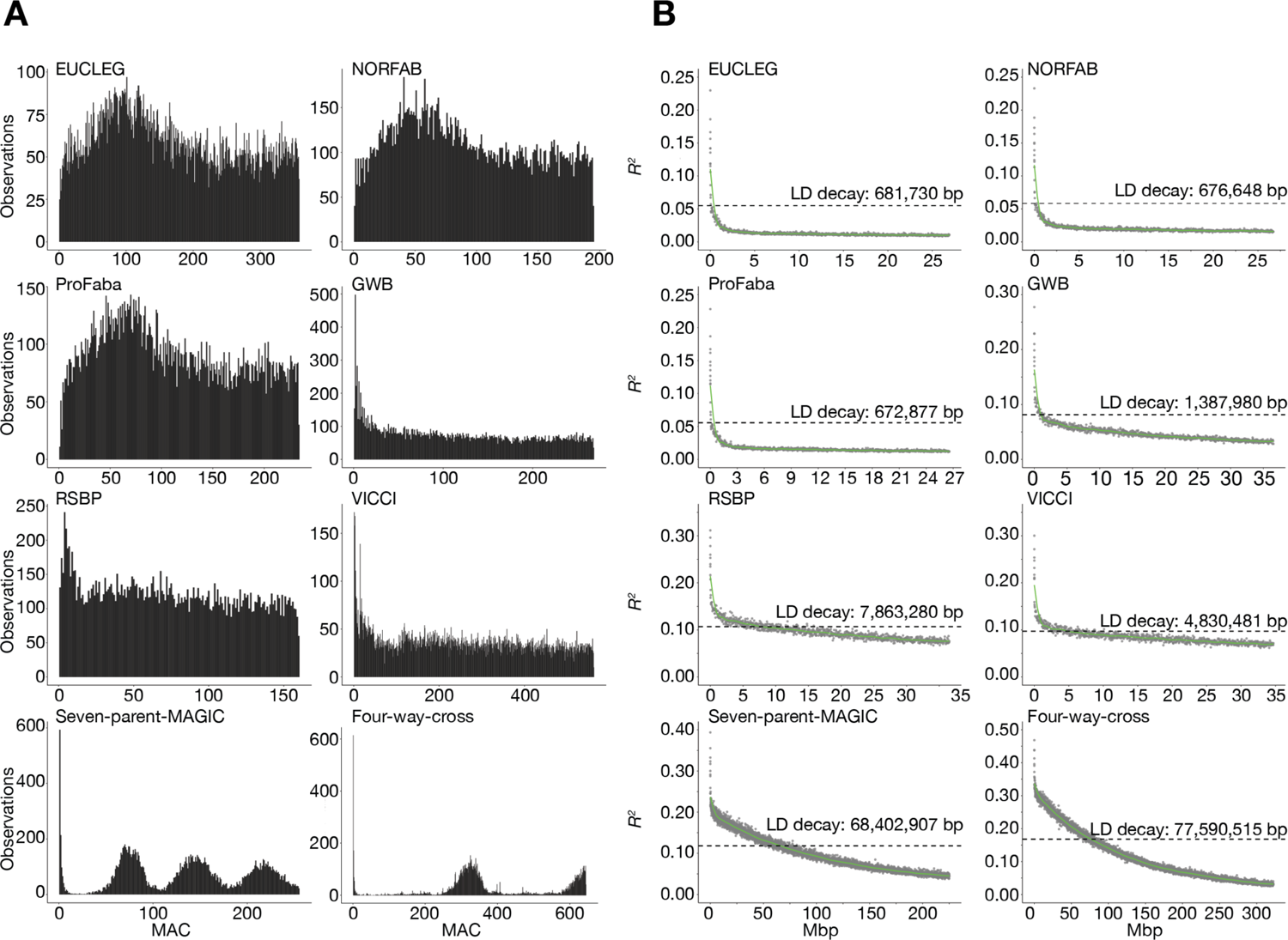
Genetic structure of individual panels. A) Folded size frequency spectrums of the eight panels show the panel-wise distribution of minor allele counts (MAC). B) Panel-wise LD decay plots. Y-axis displays the average squared correlation coefficient (*R^2^*) between markers when sorted after the average distance and binned into groups of 1000 or 5000 Seven-parent-MAGIC and Four-way-cross). For each bin, the x-axis displays the average distance in Mbp between two SNPs. The green line is the fitted loess curve with half its maximum *R^2^* indicated by the dashed line.

EUCLEG, NORFAB, and ProFaba showed similar patterns of fast LD decay with LD dropping to half of its maximum at values close to the average distance between SNP—that is, 681.7 Kbp, 676.6 Kbp, and 672.9 Kbp, respectively. LD decayed over larger distances for the GWB (1.4 Mbp), VICCI (4.8 Mbp), and RSBP (7.8 Mbp) panels consistent with the decreasing number of generations of outcrossing embodied in the respective panels. The Seven-parent-MAGIC and Four-way-cross panels showed much larger LD blocks with average decay values of 68.4 Mbp and 77.6 Mbp, respectively (**Figure 1B**).

### Key agronomic traits in the Seven-parent-MAGIC population

To get a better understanding of the genetic basis of key agronomic traits in faba bean and the associated underlying genetic regions, the Seven-parent-MAGIC panel was phenotyped for a wide range of traits during the two years of field trials at two locations in Denmark. The distribution of the obtained data can be seen in **Supplementary Figure 2**. The data collected for all traits were subjected to ANOVA, and heritabilities were calculated. In the multi-environmental models, all traits had a statistically significant contribution from the genotypic variance to the overall phenotypic variance (*p* < 0.05). Additionally, we found that the replicate variance, the environmental variance, and the GxE interaction were significant (*p* < 0.05) for all traits, except susceptibility to downy mildew as measured in percentage (**Supplementary File 4**). Seed traits showed relatively little environmental influence and high heritabilities of 0.96–0.98 (**Supplementary Figure 3, Supplementary File 4**). When including data from multiple environments, we found low heritabilities for disease resistance to chocolate spot as measured in percentage (0.21) and rust (0.28–0.30) (**Supplementary File 4**). When considering environments separately, however, heritabilities above 0.50 could be found for at least one of the environments scored for these traits (**Supplementary File 4**). Because of this and the significant GxE interactions for almost all traits, we performed GWAS for each environment separately by using the BLUEs of combined environments.

### Genome-wide association studies for key agronomic traits

GWAS was performed using FarmCPU. The Q-Q-plots associated with the GWAS results raised no concerns regarding genomic inflation, thereby revealing appropriate fitting of the models (**Supplementary Figure 4**). We identified 238 (177 unique) markers associated with statistically significant signals for the following traits: earliness of flowering; plant height; lodging; sterile tillers; seed length, width, and area; TGW; herbicide damage; and susceptibility to chocolate spot, rust, and downy mildew **(Supplementary File 5)**. Manhattan plots for multi-environmental traits—that is, susceptibility to chocolate spot, rust, and downy mildew; plant height; lodging; earliness of flowering; TGW; seed area, length, and width—are shown in **Figure 2**. Manhattan plots for the remaining traits can be seen in **Supplementary Figure 5**.

**Figure 2.**
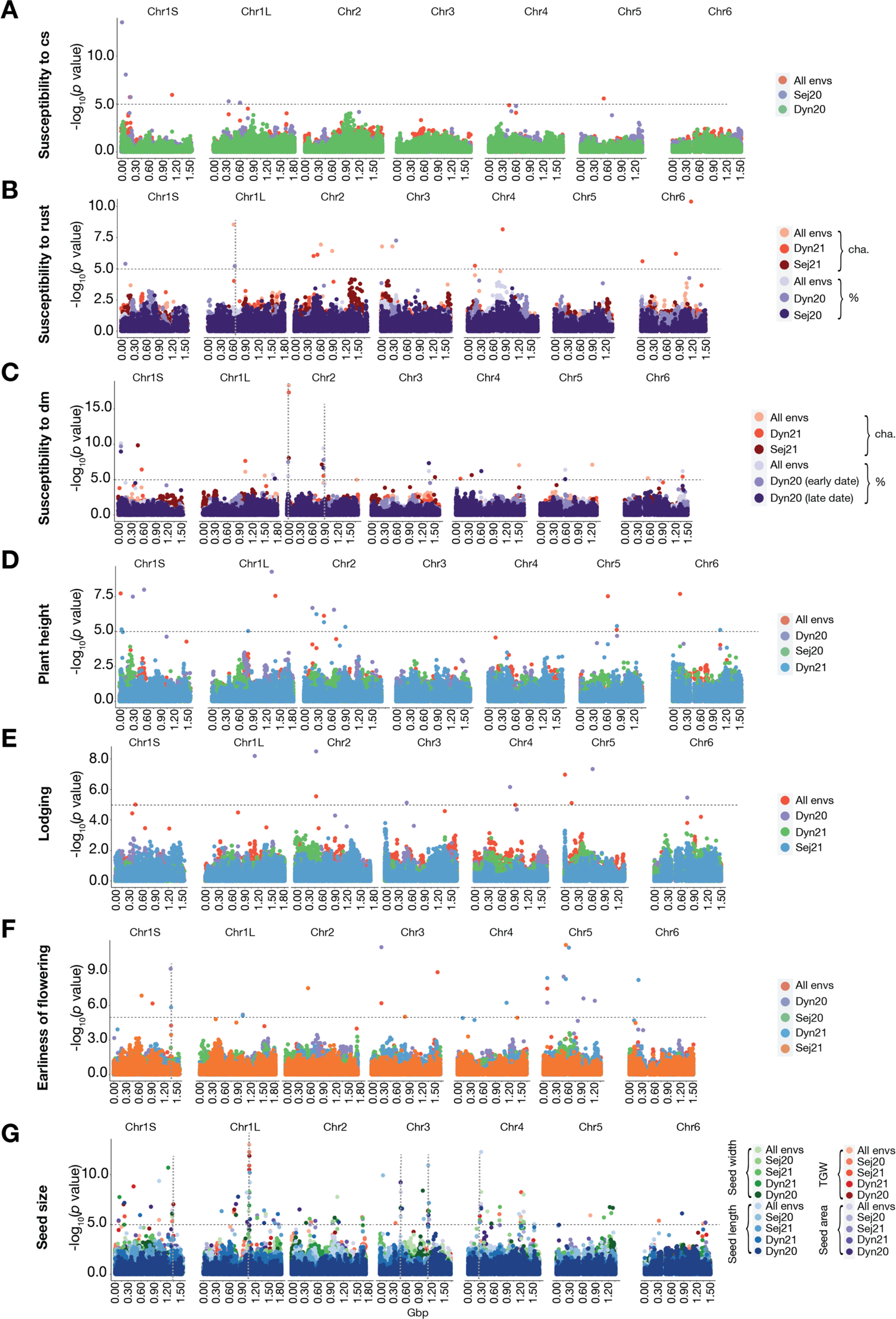
Manhattan plots of selected GWAS results in the 7-Parent-MAGIC panel. A-C) Disease susceptibility to chocolate spot (A), rust (B), and downy mildew (C). D) Plant height. E) Lodging. F) Earliness of flowering. G) Seed size traits, i.e. TGW, seed area, seed width, and seed length. The dashed horizontal line indicates the SimpleM-corrected threshold for significance. The dashed vertical lines display broad genetic regions (peaks) made up of relatively close markers associated with multiple environments and/or measurements of the same trait. Abbreviations: cha, character; cs, chocolate spot; dm, downy mildew; envs., environments.

Of the 238 marker-trait associations, 230 originate from multi-environmental traits. Of these, 65 were stable across all environments and the associated markers explained between 0.03% (TGW) and 21.8% (seed width) of the overall trait variation. Although only 10 of these markers point to major-effect QTLs (PVE(%) > 10%), they are, due to their stability, considered to report reliable trait-associated loci. In addition to single stable markers across environments, overlaying the Manhattan plots resulting from GWAS of multiple environments of the same trait allowed us to identify broad genetic regions (peaks) made up of close markers associated with multiple environments and/or measurements of the same trait. Such peak-contributing genomic regions were also considered to be highly reliable candidates in identifying stable QTLs associated with traits (**Figure 2, Supplementary File 5**).

For the disease susceptibility traits, 58 marker-trait associations were identified, of which 22 were stable across environments (chocolate spot: 3/8, rust: 5/15, downy mildew: 14/35). All stable markers had a minor effect on trait variation (PVE(%) < 10%). Broader peaks were found for susceptibility to rust at Chr1L 609,902,557–635,636,923 (∼26 Mbp) and for susceptibility to downy mildew at the following genomic locations: Chr2 26,807,439–42,451,531 (∼16 Mbp) and 839,256,282–880,296,875 (∼41 Mbp) (**Figure 2A–C, Supplementary File 5)**. For plant height, 18 marker-trait associations were significant. Six of the associations were stable across environments and individually explained up to 10.8% of all trait variance (**Figure 2D**). Lodging gave rise to ten significant associations, with four of them being stable across environments. All of the markers associated with lodging were estimated to have relatively small effects (**Figure 2E**). For earliness of flowering, 21 significant associations were identified; of these 21 associations, 4—located on Chr1S, Chr3, and Chr5—were stable across environments.

Additionally, a region at Chr1S 1,352,951,752–1,362,763,661 (∼10 Mbp) seemed to be associated with the trait in multiple environments (**Figure 2F**). A total of 123 (83 unique) significant markers were identified for traits related to seed size—that is, seed area, seed width, seed length, and TGW (**Figure 2G**). Interestingly, we identified genomic regions that were associated with multiple seed size-related traits and were stable across many environments; therefore, these can be regarded as highly reliable loci for controlling seed size. The most remarkable of these was a 26 Mbp region at Chr1L (1,049,955,413–1,075,870,570) that consists of 13 significant marker-trait associations and appears to be a very convincing candidate region for harbouring genes controlling seed size. There are 101 genes that underlie this region.

Additional stable regions associated with seed size traits were found at Chr1S 1,318,461–1,347,658,420 (∼29 Mbp); Chr3 479,473,217–484,841,073 (∼5 Mbp), Chr3 1,012,848,106–1,140,401,114 (∼128 Mbp), and Chr4 269,654,967–299,822,811 (∼30 Mbp) (**Figure 2G**).

### A panel capturing the global faba bean diversity

To investigate the genetic characteristics of the eight panels, a PCA plot was generated based on the 2,678 studied accessions (**Figure 3A**). Given the high number of accessions, the first two principal components (PCs) explained a noticeable share of the overall genetic variance (10.1%). We found that the plot showed a clear panel structure. Most obvious was the Four-way-cross accessions, which formed a tight cluster clearly separated from the remaining panels. Additionally, the GWB accessions also formed a tight cluster, suggesting that winter varieties to a high degree are genetically distinct from the spring varieties that the remaining panels primarily consisted of.

**Figure 3.**
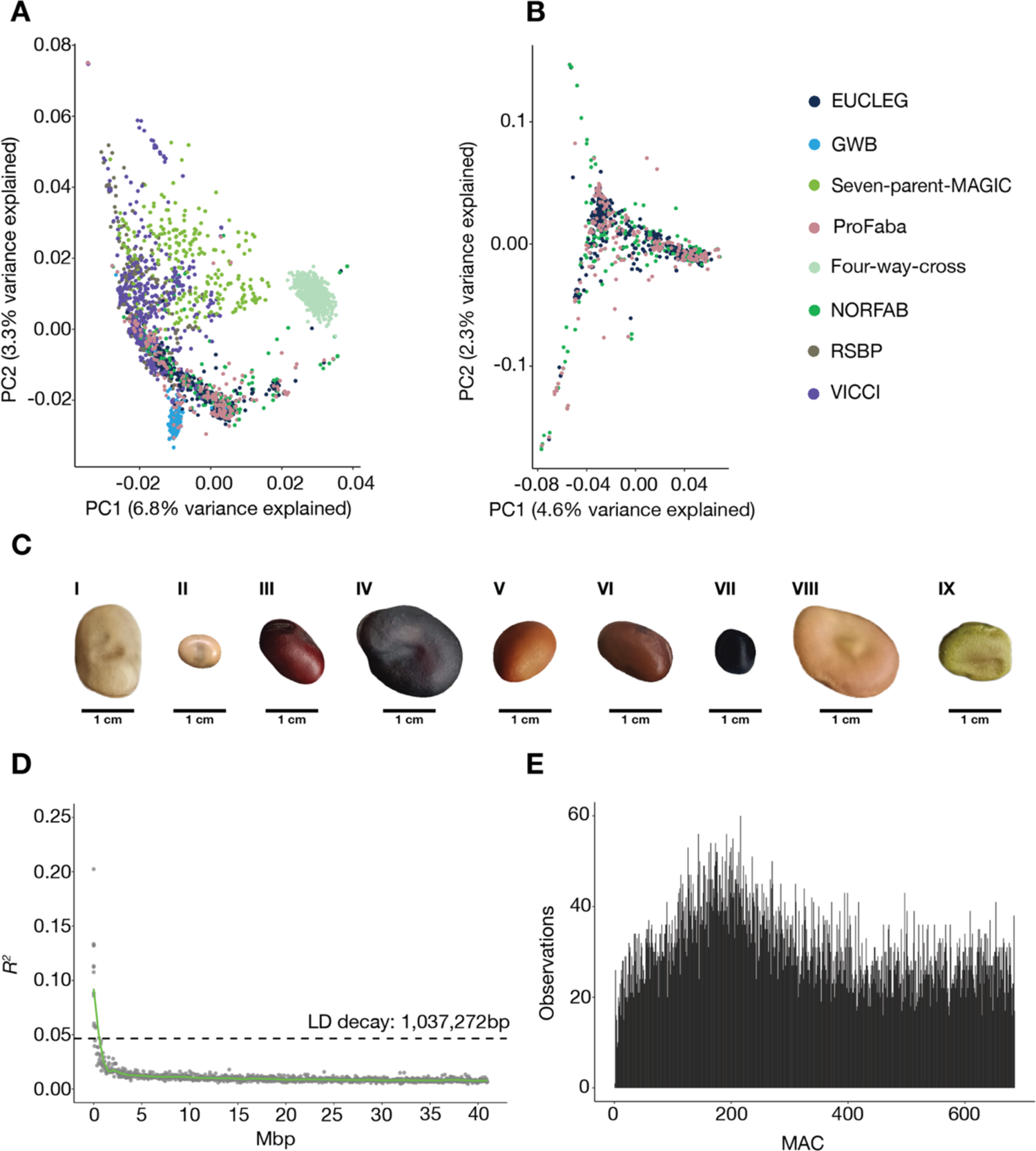
Characterisation of the diversity panel. A-B) The genetic structure of the data, as indicated by the first and second principal components and colour-coded by panel membership A) All panels, n = 2678. B) The inbred EUCLEG, ProFaba, and NORFAB diversity panels, n = 787. C) An image-based representation of the large phenotypic variation of seeds in the diversity panel. I) GPID_00080, II) EUC_VF_131, III) GPID_00162, IV) GPID_00176, V) GPID_00163, VI) GPID_00119, VII) GPID_00004), VIII) EUC_VF_272, IX) GPID_00042. D) LD decay plot for the diversity panel. Y-axis displays the average squared correlation coefficient (*R^2^*) between markers when sorted after the average distance and binned into groups of 1000. For each bin, the x-axis displays the average distance in Mbp between two SNPs. The green line is the fitted loess curve with half its maximum *R^2^* indicated by the dotted line. E) Folded site-frequency spectrum of non-monomorphic SNPs in the diversity panel. The x-axis reports the minor allele counts.

To establish a diversity panel of inbred lines, we removed the populations that were outbred (VICCI) or generated from a limited number of founders (Seven-parent-MAGIC, Four-way-cross, RSBP, GWB). This left us with 787 combined accessions from the EUCLEG, NORFAB, and ProFaba panels (**Figure 3B**). The accessions mixed well in the PCA, showing no underlying panel structure. The resulting diversity panel was then filtered for redundancy, which removed 102 samples and resulted in a large diversity panel of 685 non-redundant lines. For all subsequent analyses on the diversity panel, except the nucleotide diversity of genetic subpopulations, a 1% MAF filter was applied to the genotype data, leaving 21,116 markers.

Passport information for the diversity panel is included in **Supplementary File 1**. The lines in the diversity panel have a wide range of geographic origins representing 52 countries. In addition, they exhibit large seed variation, with seeds ranging widely in their size, colour, and morphology, as exemplified in **Figure 3C**. The genetic characteristics of the diversity panel were very similar to those of the individual panels of EUCLEG, NORFAB, and ProFaba. The average chromosomal LD decay dropped to half of its maximum at 1.0 Mbp and the folded site frequency spectrum showed a similar pattern to the MAC distributions of EUCLEG, NORFAB, and ProFaba (**Figure 1, Figure 3D+E)**.

### Population structure of the diversity panel

Running ADMIXTURE with K = 3, we found a clear correlation between the coarse geographic origin of accessions and their ancestral proportions (**Figure 4A+B)**. The correlation was not further resolved by setting K = 4 (**Supplementary Figure 6**). The coarse geographic groups presented here are as follows: “North” covers countries located in Northern and Central Europe, Canada and Russia; “South” covers countries located in Southern Europe, South America, Africa and Australia; “Middle East” covers countries located in the Middle East; and “Asia” predominantly covers countries located in Central and East Asia. Based on membership coefficients, accessions were assigned to a subpopulation (SP). We then carried out a PCA based on the genotypes, and we coloured and shaped individual points by their SP membership and geographic origin, respectively. Using both PC1 and PC2, we were able to separate accessions based on SP. PC1 distinguishes SP1 members from SP2 and SP3 members, whereas PC2 was able to further distinguish SP2 and SP3 (**Figure 4C**). To further characterise the three inferred SPs, we looked at the exact distribution of SPs per country represented in the data (**Figure 4D– F**). **Supplementary File 1** includes information on geographic origin on 406 of the lines.

**Figure 4.**
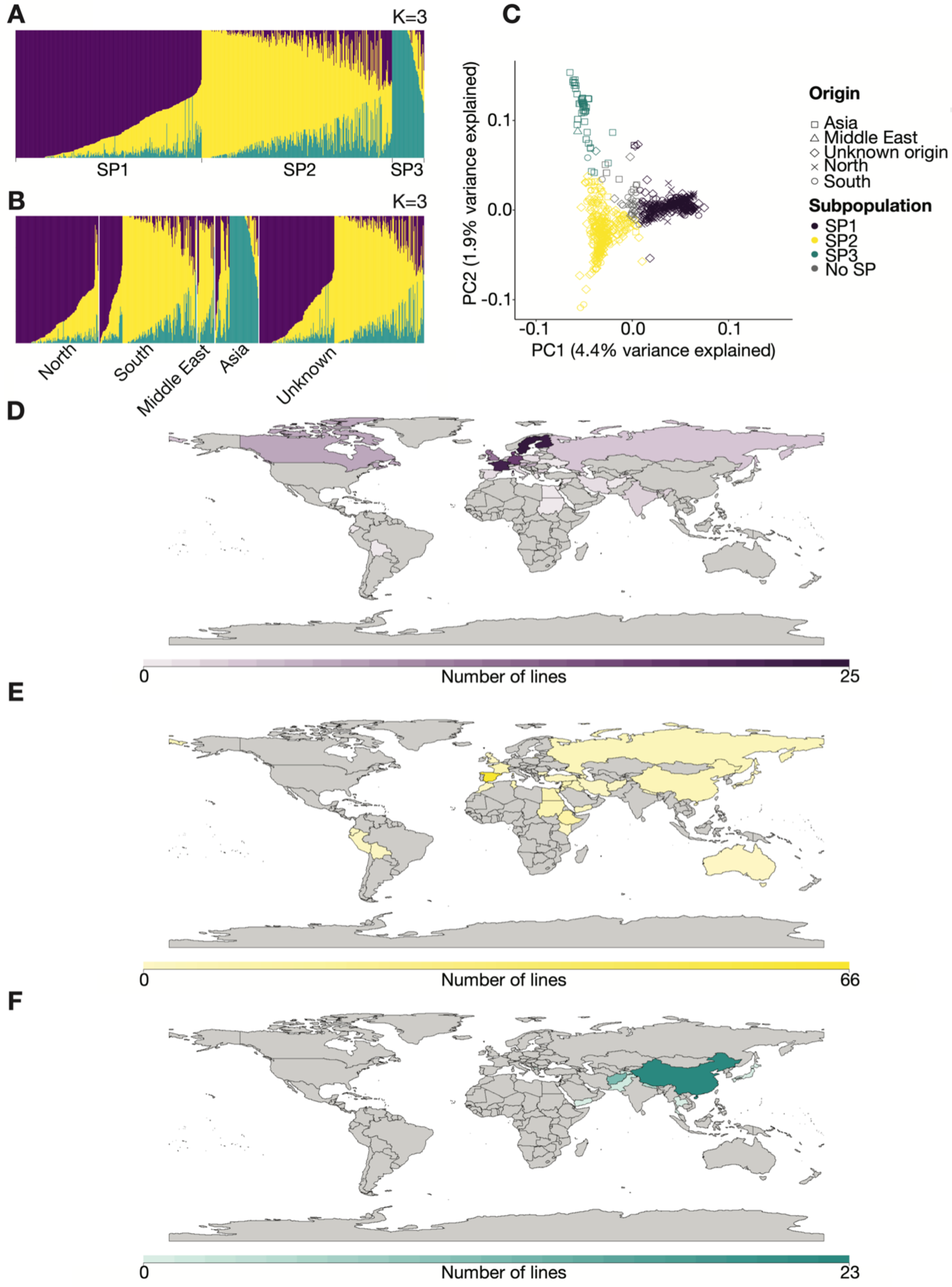
Population structure and subpopulations of the diversity panel. A-B) ADMIXTURE plots at K = 3. Each vertical bar represents a single accession coloured by its ancestry proportions. Accessions are grouped according to their subpopulation membership (A) or by their geographic origin (B). C) Principal component analysis (PCA) based on genotypes. The ADMIXTURE subpopulations at K = 3 are represented by colours and geographic origin is represented by shapes. D–F) Geographical origins of accessions belonging to SP1 (D), SP2 (E), and SP3 (F). Countries are coloured by the number of SP accessions originating from the given country, as indicated by the scale at the bottom. For simplicity, eight lines with a geographic origin in ‘Scandinavia’ are plotted in Sweden.

SP1 contains 301 accessions. Of these, 178 had a known geographic origin, and 75% of those were associated with the geographical group “North”. Among the 35 accessions associated with the geographical group “South”, 23 were French. In addition to France, the most highly represented countries/regions of origin in SP1 were Scandinavia (43), Finland (24), Germany (18), and Great Britain (12).

SP2 was made up of 304 accessions, of which 161 had a known geographic origin; the vast majority (133) was associated with the geographical group “South”. Of these accessions, 66 originate in Spain, but SP2 also includes most South American and African lines, as well as 24 Middle Eastern lines.

The smallest subgroup is SP3. It consists of 49 accessions, where the vast majority (46) have a geographic origin in Central and East Asia, predominantly China (23) and Afghanistan (12). The remaining 31 accessions were considered admixed and were therefore not assigned to any population.

### Genetic differentiation of subpopulations

The genome-wide genetic differentiation between the three subpopulations was quantified by calculating pairwise *F_ST_* values. SP2 was closely related to both SP1 and SP3, showing overall *F_ST_* values of 0.06 and 0.07, respectively. SP1 and SP3 showed the highest degree of genetic differentiation with an *F_ST_* value of 0.12 (**Table 2**). These results are consistent with the ability of the PC1 to completely separate accessions assigned to SP1 and SP3 (**Figure 4C**). AMOVA analysis of the SPs found that 5.5% of the genetic variation was due to differences between SPs, while the remaining 94.5% of the variation was found within SPs (**Table 3)**. To examine the amount of genetic diversity contained within each SP, we calculated their genome-wide nucleotide diversity (π). We found that SP3 generally showed lower nucleotide diversity (π = 0.26) than the remaining SPs (π = 0.31) (**Table 2)**. To make sure that this was not due to the low sample size of SP3 as compared to SP1 and SP2, we calculated π for 1000 subsets of 49 samples from SP1 and SP2 and used those in an FDR-based approach. We never observed a π-value as small as SP3 for the reduced samples of SP1 and SP2 (FDR = 0) (**Supplementary Figure 7**).

**Table 2.** F_ST_ analysis and nucleotide diversity of subpopulations

|  | **SP1** | **SP2** | **SP3** | **$\pi$ (nucleotide diversity)** |
| --- | --- | --- | --- | --- |
| **SP1** | 1.00 | - | - | 0.31 |
| **SP2** | 0.06 | 1.00 | - | 0.31 |
| **SP3** | 0.12 | 0.07 | 1.00 | 0.26 |

**Table 3.** AMOVA analysis

| **Source of variation** | **d.f.** | **SSD** | **MSD** | **Percentage of variation** |
| --- | --- | --- | --- | --- |
| Among populations | 2 | 482764.7 | 241382.4 | 5.5 |
| Within populations | 651 | 8367111.2 | 12852.7 | 94.5 |
| Total | 653 | 8849875.9 | 13552.6 | 100.0 |

Abbreviations. d.f., degrees of freedom; SSD, sum of squared deviation; MSD, mean squared deviation.

### Candidate loci for population divergence

To explore whether the three geographically and genetically distinct SPs are under differential selection pressures and to identify genetic regions under selection, three different methods for outlier detection were applied (**Figure 5A, Supplementary File 6**). BayeScan identified a total of 18 markers with *q*-values < 0.05, which show a substantial to decisive probability (0.89–1.00) of being under diversifying selection. The number of outliers detected by the other two methods were higher, with pcadapt identifying 339 significant outliers (*q*-value < 0.05) and Ohana finding 1,596 SNPs with a likelihood ratio ≥2. Although the relative overlap between the methods was small, five markers were identified by all methods, giving rise to a confident set of markers pointing to direct targets of diversifying selection. In total, 35 markers were considered outliers by at least two of the three methods (**Table 4**). SNPs with a distance less than the average LD decay (1 Mbp) were considered a single genomic region, meaning that the analyses identified 30 genomic regions under selection, with three of the five high-confidence markers representing a single genomic region at Chr1S 17,355,793–18,116,022 bp. To get a better understanding of the trait selection that the outlier SNPs might be involved in, we looked at how they genetically segregated between subpopulations (**Figure 5B**) and at the magnitude of their *F_ST_* signals when subpopulations were compared in a pairwise manner (**Supplementary Figure 8**). We found that the 35 selection markers showed extreme differentiation between subpopulations, as compared to 35 randomly chosen markers (**Supplementary Figure 9)**. The vast majority of outlier SNPs, including two of the five high-confidence SNPs (AX-416737096 and AX-416745027), were related to divergence of SP3 from SP1 and SP2. With the coarse geographical distinction of the SPs in mind, this clearly suggests that these markers could be associated with breeding preferences in Central and Eastern Asia (**Figure 4, Figure 5B, Supplementary Figure 8**). Interestingly, we found that the remaining three (AX-416824401, AX-416760427, AX-416791399) of the five high-confidence markers covering the 760 kbp genetic region at Chr1S were associated with the differentiation of SP1 from the remaining subpopulations. The *F_ST_* values of these markers were especially large for SP1 versus SP2 when compared to the background signal (0.52–0.71), thus reflecting what could be patterns of selection during breeding in Nordic environments (**Figure 5B, Figure 6D**). Although SP2 did not show large differentiation from either SP1 or SP3 (**Table 2**), we found one SNP on chr4 (AX-181165197) that clearly separated SP2 from both remaining SPs (**Figure 5B, Figure 6D**).

**Figure 5.**
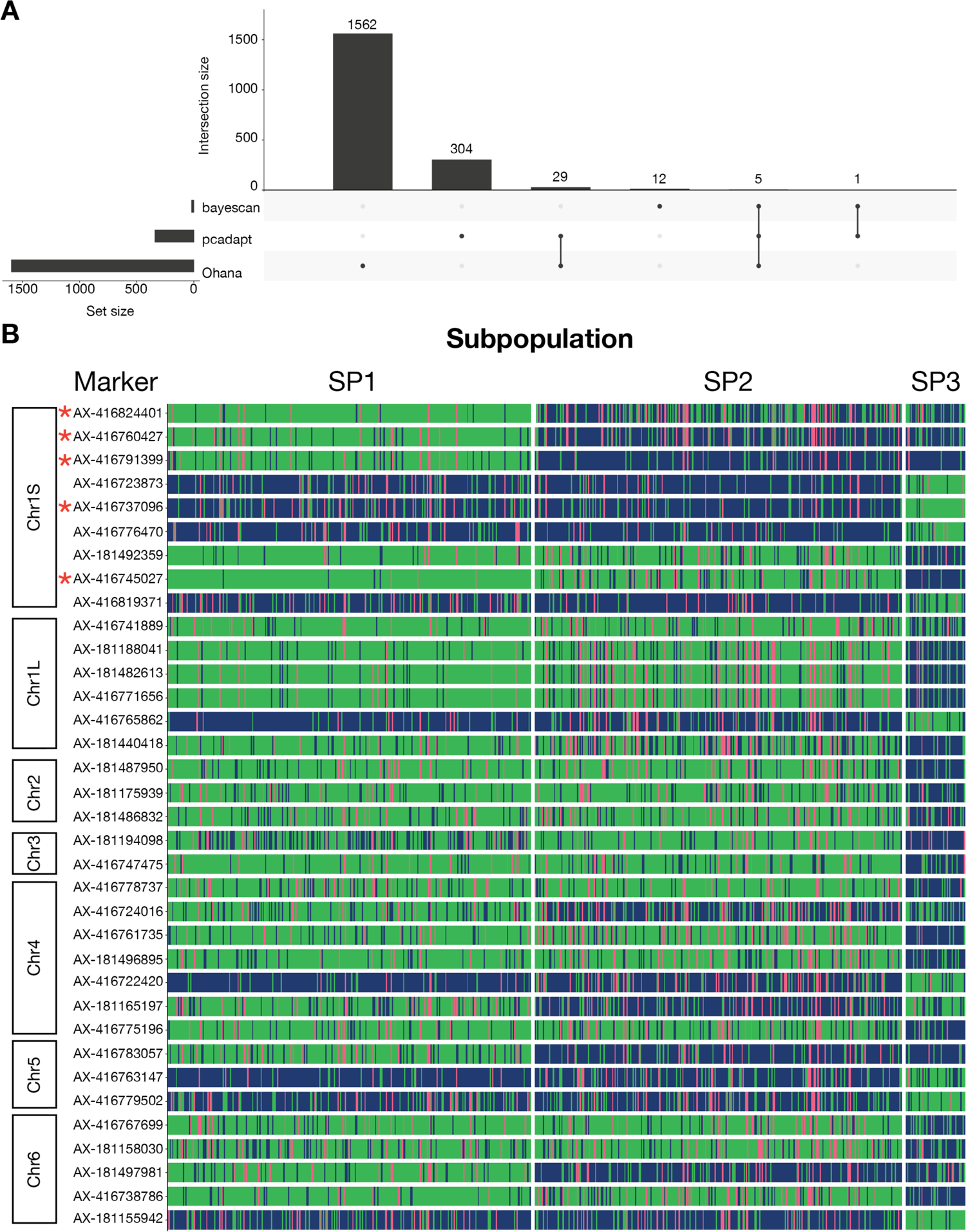
Markers under selection. A) UpSet plot of methods used for outlier detection, showing the overlapping results of BayeScan, Ohana, and pcadapt. B) Segregation of markers under selection. Each horizontal plot shows the segregation pattern of one of the 35 SNPs that shows evidence of selection. Markers are ordered according to genomic position. Each vertical line represents an accession and is coloured by genotype for a specific marker. Genotype colouring scheme is as follows: green, reference homozygote; pink, heterozygote; blue, alternative homozygote. The five high-confidence markers identified by all outlier detection methods are marked by red asterisks.

**Figure 6.**
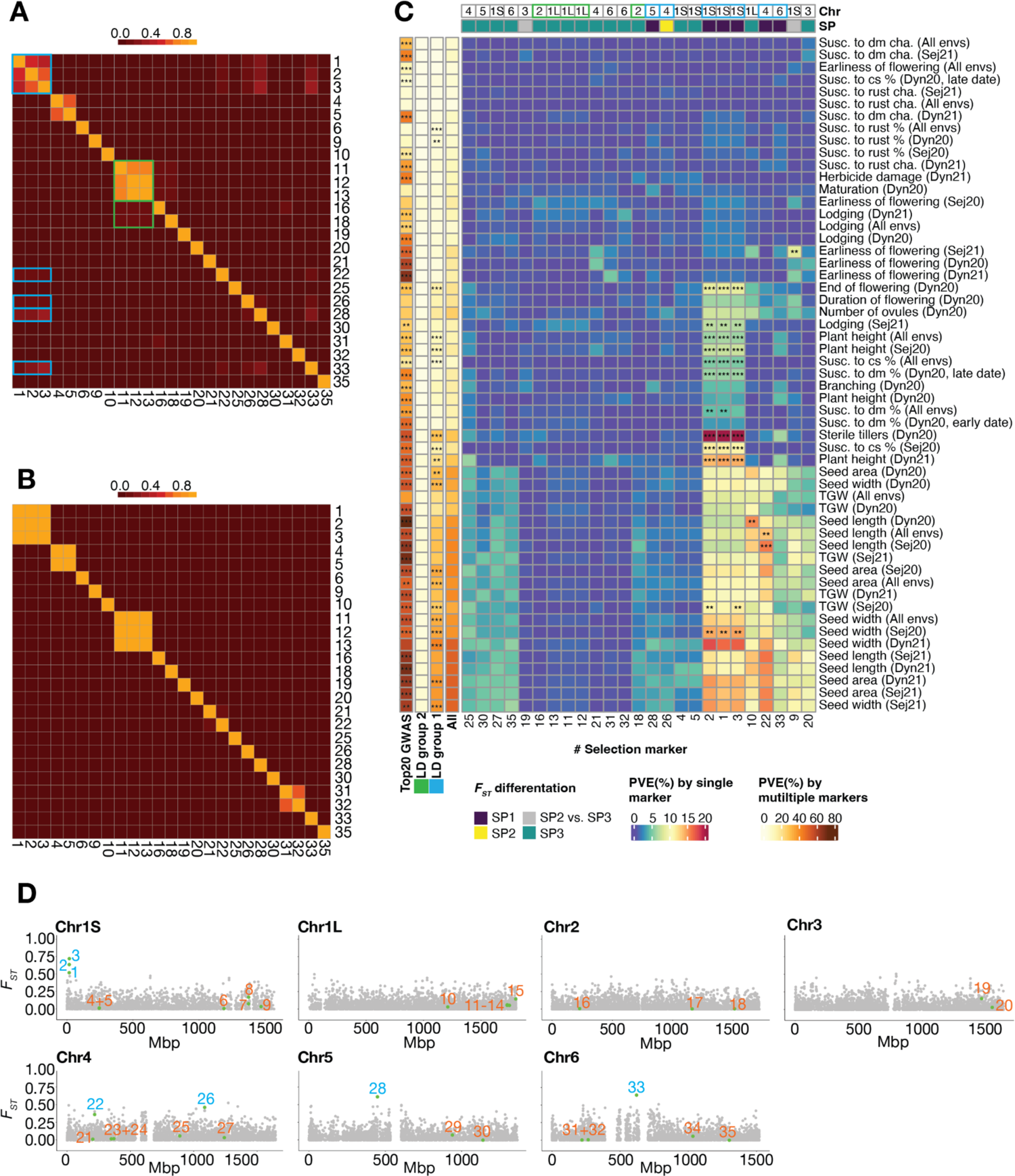
Trait variance explained by markers under selection. **A–B)** Heatmap of LD between selection markers in the diversity panel (A) or the Seven-parent-MAGIC panel (B). Markers (numerical code) are ordered according to positions in the genome. C) Proportion of variance explained (PVE) by selection markers for all traits. PVE is calculated by all selection markers individually (the large panel), all selection markers collectively (fourth column from left), top 20 most significant GWAS markers (first column from left), all markers of LD group 2 (second column from left), and all markers of LD group 1 (third column from left). At the top of the heatmap, markers are annotated by which chromosome they are located on and which SPs they differentiate: purple, differentiation of SP1 from SP2+SP3; yellow, differentiation of SP2 from SP1+SP3; teal, differentiation of SP3 from SP1+SP2; grey, differentiation of SP2 from SP3. Significance of PVE explained by different methods is calculated using an FDR-approach, where the fraction of times an obtained PVE-value was larger than what we would get from 1000 rounds of one random selected marker or different size-appropriate groups of random markers. **, 0.005 < FDR < 0.01; ***, FDR < 0.005. D) Genome-wide distribution of *F_ST_* values for SP1 vs. SP2. The *F_ST_* values of each SNP throughout a chromosome are displayed as grey dots. The green dots report the 35 SNPs under selection identified in the outlier scans. The numbers next to the green dots serves as a marker code. Selection markers in panel A-D are denoted by a numerical code: 1: AX-416824401, 2: AX-416760427, 3: AX-416791399, 4: AX-416723873, 5: AX-416737096, 6: AX-416776470, 7: AX-181492359, 8: AX-416745027, 9: AX-416819371, 10: AX-416741889, 11: AX-181188041, 12: AX-181482613, 13: AX-416771656, 14: AX-416765862, 15: AX-181440418, 16: AX-181487950, 17: AX-181175939, 18: AX-181486832, 19: AX-181194098, 20: AX-416747475, 21: AX-416778737, 22: AX-416724016, 23: AX-416761735, 24: AX-181496895, 25: AX-416722420, 26: AX-181165197, 27: AX-416775196, 28: AX-416783057, 29: AX-416763147, 30: AX-416779502, 31: AX-416767699, 32: AX-181158030, 33: AX-181497981, 34: AX-416738786, 35: AX-181155942. Markers in LD group 1 are highlighted in blue. Abbreviations: cha, character; cs, chocolate spot; dm, downy mildew; envs., environments; susc., susceptibility.

**Table 4.** Markers under selection

| SNP and genetic information |  |  |  |  | Rank |  |  |
| --- | --- | --- | --- | --- | --- | --- | --- |
| Marker | Chr | Pos | MAF | Gene Annotation* | pcadapt | Ohana | BayeScan |
| AX-416824401 | Chr1S | 17,355,793 | 0.34 | Protein NRT1 PTR FAMILY | 261 | 36 | 8 |
| AX-416760427 | Chr1S | 17,684,368 | 0.45 | Transaldolase/Fructose-6-phosphate aldolase | 120 | 540 | 3 |
| AX-416791399 | Chr1S | 18,116,022 | 0.45 | Peptidyl-prolyl cis-trans isomerase | 32 | 559 | 1 |
| AX-416723873 | Chr1S | 244,272,085 | 0.19 | alpha-L-fucosidase | 147 | 775 | 79 |
| AX-416737096 | Chr1S | 245,221,411 | 0.24 | transcription factor | 241 | 153 | 18 |
| AX-416776470 | Chr1S | 1,189,800,874 | 0.18 | Chaperone protein | 134 | 1453 | 249 |
| AX-181492359 | Chr1S | 1,375,226,525 | 0.18 | Ubiquitin carboxyl-terminal hydrolase | 140 | 1124 | 196 |
| AX-416745027 | Chr1S | 1,376,463,514 | 0.18 | Tobamovirus multiplication protein | 38 | 919 | 4 |
| AX-416819371 | Chr1S | 1,470,291,451 | 0.19 | Receptor-like cytosolic serine threonine-protein kinase | 326 | 992 | 3740 |
| AX-416741889 | Chr1L | 1,210,905,827 | 0.16 | Multiple C2 and transmembrane domain-containing protein 1 | 179 | 20357 | 15 |
| AX-181188041 | Chr1L | 1,727,444,242 | 0.20 | SNF2 family N-terminal domain | 108 | 709 | 1541 |
| AX-181482613 | Chr1L | 1,727,749,638 | 0.18 | SNF2 family N-terminal domain | 76 | 1153 | 8595 |
| AX-416771656 | Chr1L | 1,727,750,681 | 0.18 | Glucose-induced degradation protein 8 homolog | 78 | 1137 | 5621 |
| AX-416765862 | Chr1L | 1,747,192,973 | 0.19 | sphingolipid transporter spinster homologue | 166 | 871 | 6059 |
| AX-181440418 | Chr1L | 1,801,916,585 | 0.29 | Ras-related protein | 176 | 159 | 5020 |
| AX-181487950 | Chr2 | 228,005,888 | 0.21 | No annotation | 131 | 456 | 15934 |
| AX-181175939 | Chr2 | 1,160,381,636 | 0.21 | ATPase B chain family | 234 | 522 | 18858 |
| AX-181486832 | Chr2 | 1,511,910,898 | 0.21 | Involved in mitochondrial genome maintenance | 302 | 797 | 2867 |
| AX-181194098 | Chr3 | 1,465,379,679 | 0.31 | Pentatricopeptide repeat-containing protein | 211 | 1039 | 16108 |
| AX-416747475 | Chr3 | 1,549,101,456 | 0.18 | phosphatidylglycerol acyl-chain remodelling | 328 | 1541 | 9514 |
| AX-416778737 | Chr4 | 203,422,818 | 0.21 | Brefeldin A-inhibited guanine nucleotide-exchange protein | 254 | 429 | 7697 |
| AX-416724016 | Chr4 | 218,146,604 | 0.46 | GPI mannosyltransferase | 285 | 441 | 11366 |
| AX-416761735 | Chr4 | 349,618,738 | 0.22 | TIFY 10B-like | 174 | 232 | 12567 |
| AX-181496895 | Chr4 | 370,017,390 | 0.21 | Protein of unknown function (DUF1296) | 297 | 442 | 1259 |
| AX-416722420 | Chr4 | 889,829,096 | 0.21 | Telomere repeat-binding factor | 251 | 1423 | 10805 |
| AX-181165197 | Chr4 | 1,086,134,085 | 0.49 | Aldehyde dehydrogenase family | 307 | 536 | 20320 |
| AX-416775196 | Chr4 | 1,241,304,815 | 0.24 | Ubiquitin carboxyl-terminal hydrolase | 259 | 114 | 9265 |
| AX-416783057 | Chr5 | 448,641,887 | 0.44 | Pentatricopeptide repeat-containing protein | 191 | 590 | 9273 |
| AX-416763147 | Chr5 | 939,720,483 | 0.19 | Nuclear transcription factor Y subunit | 249 | 839 | 11704 |
| AX-416779502 | Chr5 | 1,140,044,946 | 0.29 | pectinesterase | 291 | 10 | 16148 |
| AX-416767699 | Chr6 | 219,883,438 | 0.18 | E3 SUMO-protein ligase | 198 | 1558 | 19473 |
| AX-181158030 | Chr6 | 265,567,017 | 0.27 | Copper transporter | 336 | 1283 | 9116 |
| AX-181497981 | Chr6 | 620,182,068 | 0.50 | zinc-RING finger domain | 151 | 740 | 15494 |
| AX-416738786 | Chr6 | 1,033,478,650 | 0.21 | Cytochrome c biogenesis protein | 220 | 1191 | 20040 |
| AX-181155942 | Chr6 | 1,300,142,339 | 0.26 | Ribosomal protein S1-like RNA-binding domain | 288 | 43 | 8995 |

\* Annotation of closest gene if the marker is intergenic

### Candidate traits under selection

To get a better understanding of the selection markers and how they have been important in the global selection during breeding of faba bean, we investigated their pairwise LD in the diversity panel (**Figure 6A**). We then compared the observed patterns with the pairwise LD in the Seven-parent-MAGIC panel (**Figure 6B**) where we were able to identify broad genetic regions associated with traits of interest (**Figure 2 + Supplementary Figure 5)**. As allele frequencies were different between the two panels, only 25 out of the 35 selection markers were included in the analyses. We found that all markers showing segregation between Northern and Southern accessions—that is, between SP1 versus SP2+SP3 or SP2 versus SP1+SP3—showed patterns of LD in the diversity panel (blue boxes, **Figure 6A**). This group of markers (referred to as ‘LD group 1’) consists of the three adjacent high-confidence markers at Chr1S, which due to their physical proximity are fully linked in the Seven-parent-MAGIC panel, as well as the four remaining markers, which give rise to unusual patterns of long-range LD, since they are located on Chr4, Chr5, and Chr6 and consequently lose their LD in the Seven-parent-MAGIC (**Figure 6A+B**). In addition, we found another group of selection markers showing long-range LD in the diversity panel (green boxes, **Figure 6A**). This group, referred to as LD group 2, was associated with the differentiation of Asian lines—that is, SP3 versus SP1+SP2 (**Figure 5 + 6A**). After recombination in the Seven-parent-MAGIC panel, the adjacent markers of LD group 2 showed full LD, whereas long-range LD was broken down.

To investigate possible links between the genomic regions under selection and specific traits, we resorted to the Seven-parent-MAGIC panel. Because of the long range of LD in this panel, QTL positions could not be determined with great accuracy. This means that the co-localisation of selection markers and QTLs had to consider very large genetic regions, making an overlap by chance very likely. For this reason, we took a different approach. For each trait subjected to GWAS in the Seven-parent-MAGIC panel, we calculated the proportion of phenotypic variance explained by each marker and LD groups under selection in the diversity panel (**Figure 6C**). For comparison, we tested how much of trait variation could be explained by the collective effort of all 25 selection markers and top 20 most significant GWAS markers. We used top 20 markers because the 25 selection markers, when considering LD in the Seven-parent-MAGIC population, behaves as 20 markers (**Figure 6B).**

Most of the selection markers were not found to explain any statistically significant proportion of variance for any of the traits. However, four of the selection markers in LD group 1 were able to individually explain a statistically significant proportion of the variance of one or more traits. Most remarkable were the three adjacent markers at Chr1S covering a 760 kbp genetic region, which explained a statistically significant proportion of the phenotypic variance of traits related to seed size, plant height, end of flowering, lodging, sterile tillers, and disease resistance to downy mildew and chocolate spot (**Figure 6C**). These markers were among the most differentiated for SP1 and SP2 (**Figure 6D and Supplementary Figure 8**). The fourth marker was located at Chr4 and explained a significant proportion of trait variance for seed length (**Figure 6C**). Expanding the single markers to the entire LD group 1, the list grew with the following additional traits: susceptibility to rust and several additional traits related to seed size traits. All traits that could be explained by selection markers were better explained by top GWAS markers that generally explained a large proportion of the overall trait variance. An exception to this observation is susceptibility to rust, where top GWAS markers explained no significant part of the trait variation, while LD group 1 markers did (**Figure 6C**).

To better disentangle the traits significantly explained by the selection markers associated with SP1 versus SP2 differentiation (that is, LD group 1), we looked at correlations between genetic values of the traits (as used in GWAS) (**Supplementary Figure 10, Supplementary File 7**). Traits related to seed size were correlated with the following four types of traits that showed no correlations with each other: end of flowering (negative), susceptibility to rust (negative), sterile tillers (positive), and lodging (positive). Additionally, susceptibility to chocolate spot had a positive correlation with sterile tillers and lodging. Plant height and susceptibility to downy mildew show no correlation with any of the other measured traits (**Supplementary Figure 10, Supplementary File 7**). From the results it seems likely that multiple traits might have been co-selected during breeding for different market types or environments.

## Discussion

### Characterisation of individual panels

Using 21,345 genome-wide high-quality SNPs, we performed genetic analyses on a large collection of faba bean germplasm. Few other studies on faba bean diversity have combined such a large number of geographically diverse accessions and high-quality SNP markers to study genetic diversity. Our results of the eight panels studied revealed genetic diversity reflecting the underlying panel structure. Most remarkable was GWB, a population derived from 11 winter-type founders, which was found to be clearly genetically distinguishable from the remaining panels. As the remaining panels predominantly consisted of spring-type germplasms, this suggests that winter-type and spring-type cultivars are highly genetically distinct. A similar distinction between winter and spring-types has been described in Chinese germplasm (Zong et al. 2009; Wang et al. 2012).

The site frequency spectrum of the diversity panels revealed a relatively uniform distribution with a slight overrepresentation of markers with intermediate allele frequencies (∼0.1–0.3). This pattern is expected because of the ascertainment bias of the utilised Axiom SNP array. The ascertainment bias is caused by the SNP discovery process using only 12 individuals for the discovery panel, with preference given to alleles of intermediate frequency with a high polymorphism information content (Albrechtsen et al. 2010).

The nucleotide diversity of the individual panels ranged from 0.26 to 0.32. As expected, the lowest genetic diversity was found for populations established from a limited number of founders, with the Four-way-cross being the most extreme. The highest nucleotide diversities were found for the diversity panels (π = 0.32) and the outbreeding population (VICCI, π = 0.30). The nucleotide diversity in the combined diversity panel (n = 685) was 0.31. These values are similar to those reported using SNP data in inbred panels of maize, where values between 0.27 and 0.39 have been estimated (Hamblin et al. 2007; Lu et al. 2009; Van Inghelandt et al. 2010; Yang et al. 2011; Bouchet et al. 2013; Shu et al. 2021). The highest genetic diversity (0.39) stems from a population of 527 inbred maize lines with very broad origins (Yang et al. 2011).

### Mapping of agronomic traits

Few studies have been performed to identify QTLs of agronomically important traits in faba bean (Khazaei et al. 2021). Furthermore, the few published studies have typically relied on biparental populations, limiting the amount of genetic variation studied as compared to a MAGIC population. Here, we use GWAS to identify 238 significant marker-trait associations linked to 12 agronomic important traits. Of these marker-trait associations, 65 (27%) were found to be stable across multiple environments, thereby pointing to high-confidence candidate regions for harbouring genes associated with plant height, stem lodging, earliness of flowering, seed size, and resistance to chocolate spot, downy mildew, and rust. Furthermore, all traits scored in multiple environments gave rise to stable QTLs. Among these we found major QTLs (PVE > 10%) for TGW (11.0–16.8%), seed width (13.0–21.8%), seed length (16.4–19.2%), seed area (13.5–14.3%), and plant height (10.8%). As these QTLs have a major effect and have been associated with 3–4 different Danish environments, they provide clear targets for integration in future breeding programs.

Earlier studies have identified several stable QTLs associated with seed size on chromosomes 2, 4, 5, and 6 in faba bean (Khazaei et al. 2014; Ávila 2017). Here, we found traits related to seed size to be highly polygenic with stable signals on all chromosomes. Especially striking is the tall peak identified at chromosome 1L position 1,049,955,413–1,075,870,570 bp, which consists of markers significantly associated with multiple traits related to seed size (TGW, area, length, width) scored in multiple environments. Markers here explained between 0.1 to 15.8% of phenotypic variation.

Plant height is another important trait related to the final crop yield of faba bean. Previous studies have performed QTL mapping of plant height but have not identified any stable QTLs across environments (Ávila 2017). In this study, we detected six QTLs that were stable across three Danish environments for plant height. The stable markers individually explained between 0.2 and 10.8% of phenotypic variation.

The faba bean genetics of flowering time has been explored earlier, as a major stable QTL has been found on chromosome 5 (Cruz-Izquierdo et al. 2012; Catt et al. 2017). Interestingly, the region was not only found to have a large effect on the trait but is also highly conserved in multiple legumes, including *Lotus japonicus* (Gondo et al. 2007), *Medicago truncatula* (Pierre et al. 2008), chickpea (Cobos et al. 2009), narrow-leafed lupin (Nelson et al. 2006), and alfalfa (Robins 2007). Although we identified a stable QTL of flowering at chromosome 5, none of our stable flowering-related QTLs were estimated to explain a large proportion (>10%) of the trait variation. On the contrary, our findings suggest a relatively polygenic nature of flowering, with multiple QTLs specific to environments.

Stable QTLs for number of ovules and branching (number of branches with flower) has previously been reported on chromosomes 3 and 6, respectively (Ávila 2017), but here we report no QTLs related to these traits.

One of the main threats for the global production of faba bean is foliar diseases such as rust (caused by *Uromyces viciae-fabae*), chocolate spot (caused by *Botrytis fabae*), and downy mildew (caused by *Peronospora viciae*). Due to environmental and economic reasons, breeding for disease resistance is preferred over treating crops with fungicides (Stoddard et al. 2010). Still, the genetic basis of faba bean disease resistance is to a large extent unknown.

Here, we identified several genomic regions associated with resistance towards all three fungal diseases. We especially obtained many stable marker-trait associations (14) for downy mildew, where we found very strong peaks on chromosome 2, at positions 26,807,439-42,451,531 bp and 839,256,282-880,296,875. This is of great interest, as no QTLs of this trait have, to our knowledge, been published for faba bean. Similar to recent studies, we found that chromosome 1 harbours QTLs associated with resistance to chocolate spot (Gela et al. 2022). For rust resistance, we found five stable markers located at chromosomes 1L, 2, and 3. Two genes associated with rust resistance in faba bean, *Uvf2* and *Uvf3*, have successfully been identified and mapped to chromosomes 3 and 5, respectively, using KASP markers (Ijaz et al. 2021). By mapping the KASP markers to our reference genome, we did not observe any overlap between the genetic regions associated with *Uvf2* and *Uvf3* and our peaks for rust resistance.

Although we detect many high-confidence QTLs associated with key agronomic traits, the low resolution in the Seven-parent-MAGIC population complicates the search for underlying candidate genes. As compared to the diversity panels, where almost no LD were detected between neighbouring SNPs, larger regions of LD were observed for the Seven-parent-MAGIC population. For this reason, the GWAS is expected to cover close to all genome-wide QTLs.

However, this is accompanied by a poor mapping resolution when it comes to identifying genes associated with traits of interest. As the average genome-wide distance between annotated genes is 307,734 bp and the LD-decay in the population is ∼68 Mbp, each marker-trait association is expected to report a region representing hundreds of genes. With this in mind, the presented GWAS results are useful in associating traits with mapped but relatively broad underlying genetic regions. For this reason, we suggest that future studies take advantage of the diversity panel for fine-mapping of the QTLs.

### Faba bean diversity and genetic differentiation

With a long history of cultivation and widespread adaptation, faba bean provides excellent material for studying global genetic diversity. In order to understand the genetic differentiation related to different geographic regions, we divided the diversity panels into three subpopulations with different coarse geographic origins: SP1, consisting of germplasm originating mostly from Northern and Central Europe but also including all Canadian lines; SP2, which mostly consists of Spanish germplasm but also includes African, South American, and Middle Eastern varieties; and SP3, which has a narrower geographic origin, mostly consisting of Central and East countries of Asia, predominantly China and Afghanistan.

Consistent with previous studies, our analyses revealed that the genetic diversity of faba beans was highly associated with geographical origin (Kaur et al. 2014b; Wang et al. 2012; Zong et al. 2010; El-Esawi 2017). Outcomes of our PCA and *F_ST_* studies identified the Northern accessions (SP1) and Central and East Asian accessions (SP3) as genetically distinct subpopulations with Southern accessions (SP2) located in between. This is also demonstrated by very few accessions showing a high degree of admixture between SP1 and SP3 and close to no geographical overlap between SP1 and SP3. Geographically, our findings fit well with the proposed routes of migration for faba bean cultivation, suggesting that different routes radiated from the Middle East (SP2) further east (SP3) and to Europe (SP1 and SP2), either via Africa (SP2) or Southern Europe (SP2) (Cubero 1974).

Consistent with our findings, previous studies have reported that Asian, or specifically Chinese, germplasm is highly distinct from other germplasm (Kaur et al. 2014b; Wang et al. 2012; Zeid et al. 2003). However, published articles aiming for further distinction between accessions from the non-Asian geographical regions are indecisive. Our findings agree with those of Zeid et al. (2003), who reported a close genetic relationship between Northern African lines and South European lines, which support the observed grouping of African and Southern European lines in SP2. Furthermore, Zong et al. (2010) reported genetic support of a subdivision of European lines into those originating from Spain and those from Northern European. Other studies, however, have found that germplasm from both Southern and Northern Europe cluster together and are genetically distinct to the group formed by Asian and Africa germplasm (Göl et al. 2017), distributing our SP2 into SP1 and SP3.

The level of genetic diversity was lowest for SP3, which includes most of the Central and East Asian accessions. This is in contrast to the findings published by Zong et al. (2009), where Asian lines (excluding Chinese) showed higher genetic diversity than either the African or European lines. As our findings did not seem to be a direct consequence of the low sample size of SP3 (n = 49), we speculate that it might be a consequence of SP3 mostly representing two countries (China and Afghanistan), thereby representing what might be expected to be a low effective population size compared to the remaining subpopulations.

AMOVA results revealed a higher genetic diversity within populations than between the three subpopulations. This is in agreement with what has earlier been found for faba bean (Göl et al. 2017; Wang et al. 2012; Oliveira et al. 2016). In our findings, the low degree of genetic variability observed between populations is most likely both a result of overlapping geographical regions of SP2 and the remaining SPs, as well as an indication of global exchange of germplasm. The high degree of within-population variability is most likely due to the reproductive nature of faba bean, which is partially outcrossing (Göl et al. 2017; Brünjes and Link 2021).

### Signatures of selection

Of the total markers, 35 (0.2%) were identified to be under selection by at least two of the three outlier-detection methods. In general, there was low agreement between the results of the different methods, most likely due to the different assumptions and estimation methods of the models. This helped us limit the selection signatures to a few highly confident markers that show strong differentiation between the different subpopulations. Most (26) of these markers were associated with differentiation of SP3 from SP1 and SP2, whereas only 6 markers (from four genetic regions) were associated with SP1 differentiation from SP2 and SP3. This further supports the differentiation of Northern (SP1) and Asian germplasm (SP3), with the Southern germplasm (SP2) being located somewhere in between. Especially interesting were five selection markers that were identified by all three methods. These markers, representing three regions at chromosome 1S (approximately at 17.4–18.1 Mbp, 245.2 Mbp, and 1376.5 Mbp), show very strong selection signatures and have very likely played an important role in the geographical differentiation of faba bean.

To couple the selection signatures with their associated traits, we took advantage of the Seven-parent-MAGIC panel, where we tested the amount of trait variance that markers under selections could explain compared to random markers. Interestingly, we mainly found selection markers associated with the differentiation of Northern (SP1) versus Southern (SP2) germplasm to explain a significant proportion of trait variances. With a key influence of the strongly differentiated region at Chr1S position 17.4-18.1 Mbp, the selection signatures of Northern and Southern accessions explained variance related to disease resistance, end of flowering, seed size, plant height, and lodging. This is in line with studies of selection in other domesticated crops such as chickpea (Varshney et al. 2019), soybean (Saleem et al. 2021), and maize (Bouchet et al. 2013), which found that genes underlying selection signatures are often associated with flowering or disease resistance.

Our results indicate that one or more of these traits could have played a role in selection for different market types or climatic conditions. Because of the large extent of LD in the Seven-parent-MAGIC panel, however, we are not able to pinpoint specific causal trait(s) at this stage. With comprehensive phenotyping, the better mapping resolution of the diversity panel could help to clarify this question in future studies.

## Supporting information

Supplementary File 1: Passport information on individual accessions

Supplementary File 2: SNP markers and their positions

Supplementary File 3: Trait descriptions

Supplementary File 4: ANOVA results of the Seven-parent-MAGIC population

Supplementary File 5: Significant marker-trait associations identified by GWAS

Supplementary File 6: Individual results of all three methods of outlier detection

Supplementary File 7: Correlations between genetic values of all traits

Supplementary File 8: Seven-Parent MAGIC phenotypes used for GWAS

Supplementary File 9: Raw Phenotype Scores from Field Trials

## Acknowledgements

The authors would like to thank technician Svenja Wiedenroth from Göttingen University for providing us with photos of seeds, and Taylor FitzGerald from Aarhus University for constructive criticism of the manuscript.

## Statements and declarations

### Author contributions

Conceptualization, S.U.A., C.K.S., D.M.O.; Methodology, C.K.S.; Software, C.K.S., T.R., J.K.; Validation, C.K.S, S.U.A.; Formal Analysis, C.K.S., T.R., J.K., W.E., D.A., L.I.F.; Investigation, L.K.N., A.S., J.K., A.Wa., N.G., A.Wi., V.T.; Resources, S.U.A., L.K.N., A.S., J.K., S.A., H.K., W.L, A.M.T., D.M.O., A.Wa.; Data Curation, C.K.S., T.R., W.E.; Writing – Original Draft, C.K.S.; Writing – Review & Editing, S.U.A., C.K.S.; Visualization, C.K.S.; Supervision, S.U.A., L.J., D.M.O.; Project Administration, S.U.A.; D.M.O., A.M.T., S.A., W.L.; Funding Acquisition, S.U.A., D.M.O., A.M.T., S.A., W.L., A.Wa., J.S.

### Data availability

Data supporting the findings are available within the paper and its Supplementary Information Files. Genotype data is available at: https://figshare.com/s/a30c37481c6c8f6626e8

### Code availability

Scripts used for data analyses and plotting are provided on github: https://github.com/cks2903/DiversityStudiesFabaBean2021

### Supplementary Figures

**Supplementary Figure 1.**
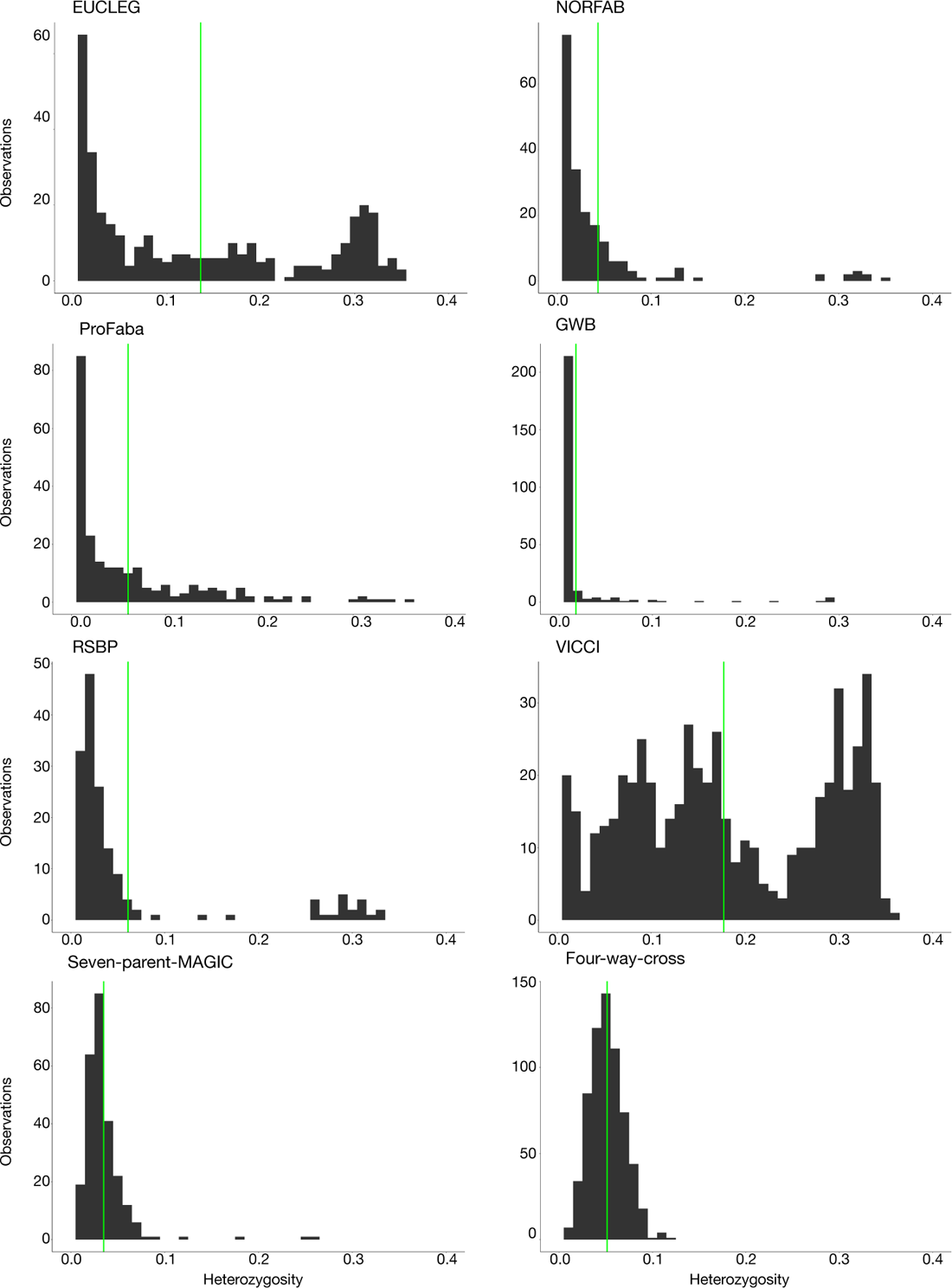
Heterozygosity of panels. Histograms showing the heterozygosity distribution within each panel. Green lines indicate the panel-wise average heterozygosity.

**Supplementary Figure 2.**
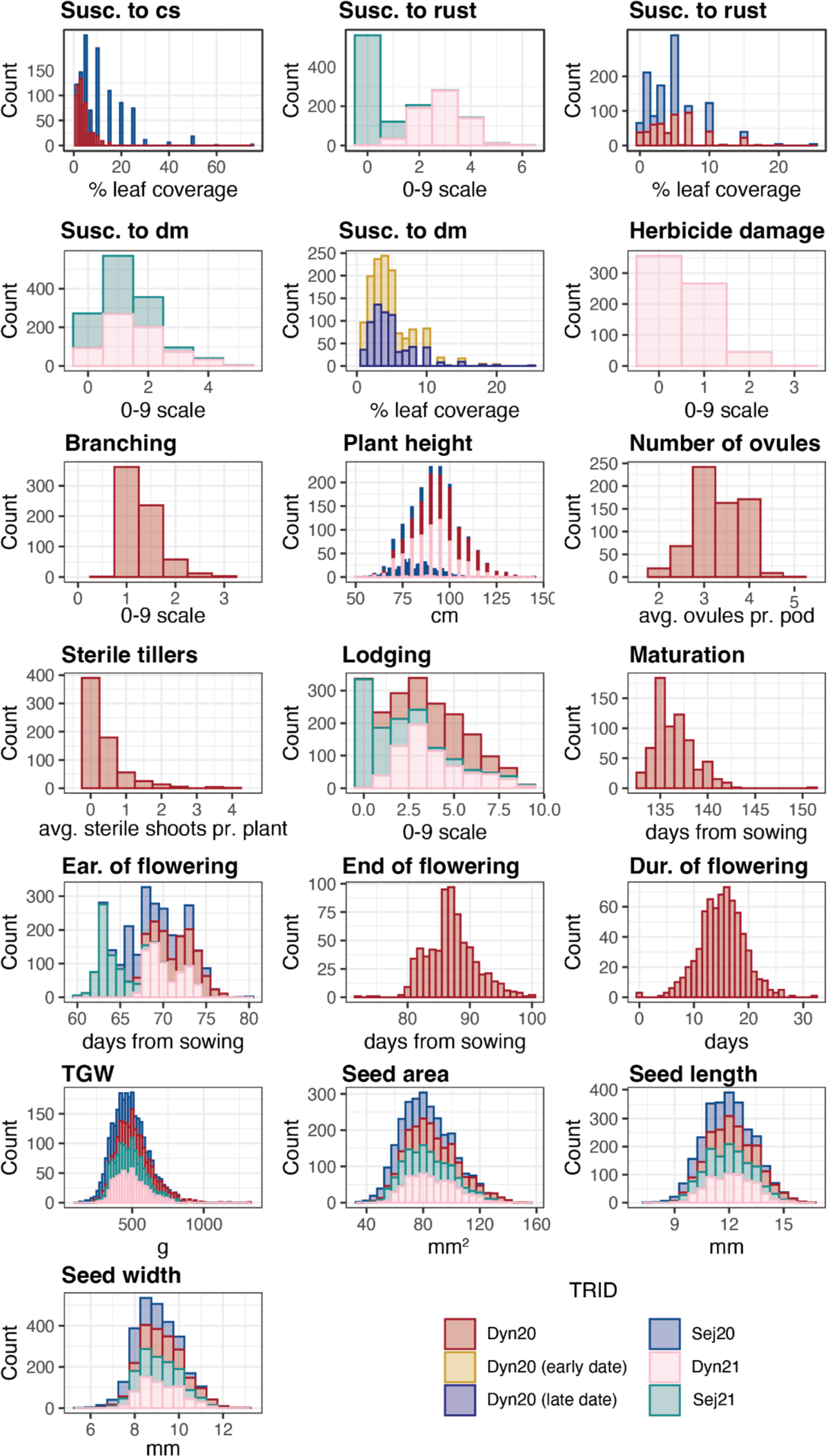
Histograms for GWAS traits. The distributions of raw phenotypes are plotted for all traits and environments. Abbreviations: cs, chocolate spot; dm, downy mildew; dur, duration; ear, earliness; susc., susceptibility.

**Supplementary Figure 3.**
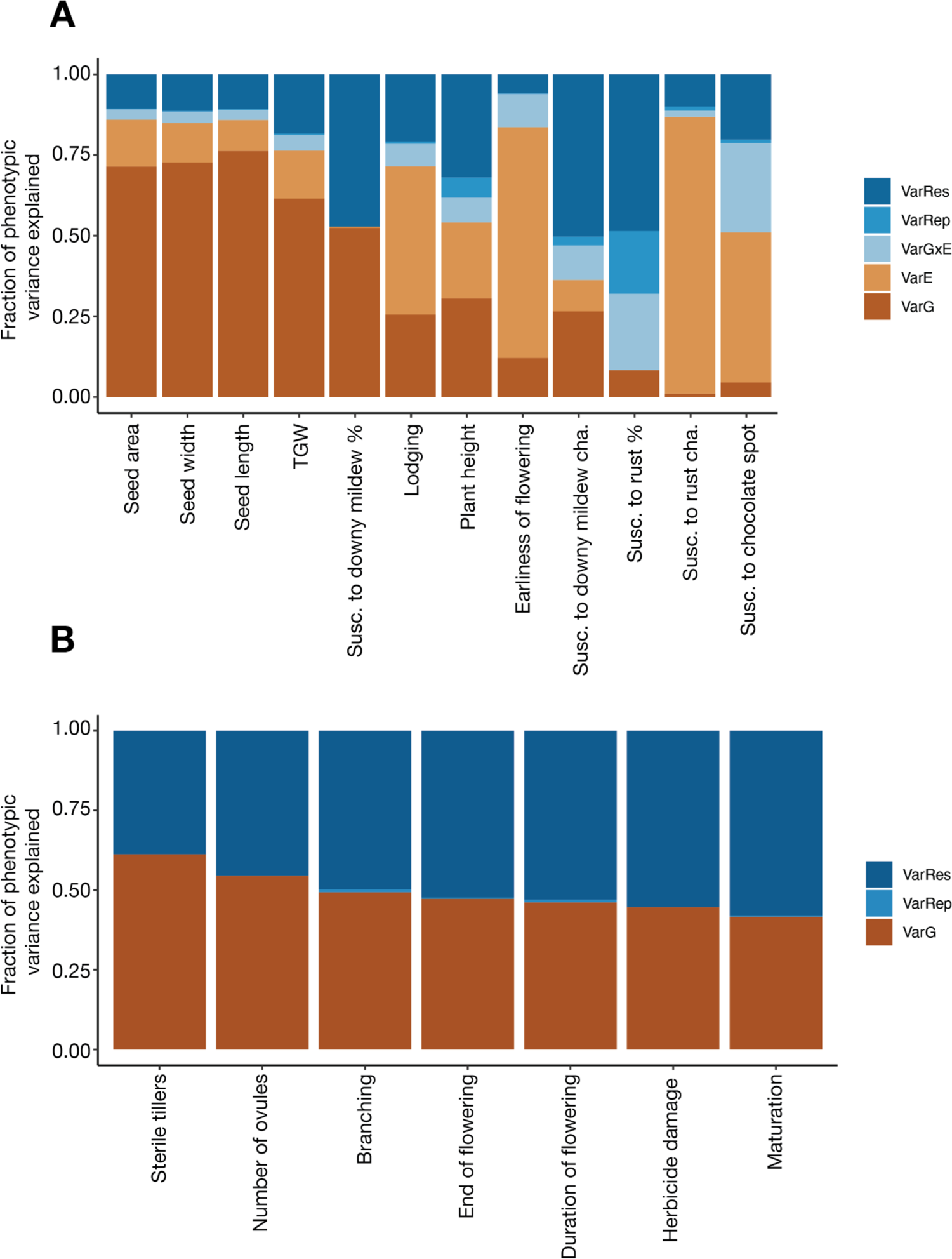
Trait variance. Proportion of phenotypic variance of MAGIC traits explained by residual variance (VarRes), replication variance (VarRep), genotype x environment variance (VarGxE), environmental variance (VarE), and genetic variance (VarG). A) All traits scored in multi-environmental field trials. B) Traits scored in one environment only. Abbreviations: cha, character; susc., susceptibility.

**Supplementary Figure 4.**
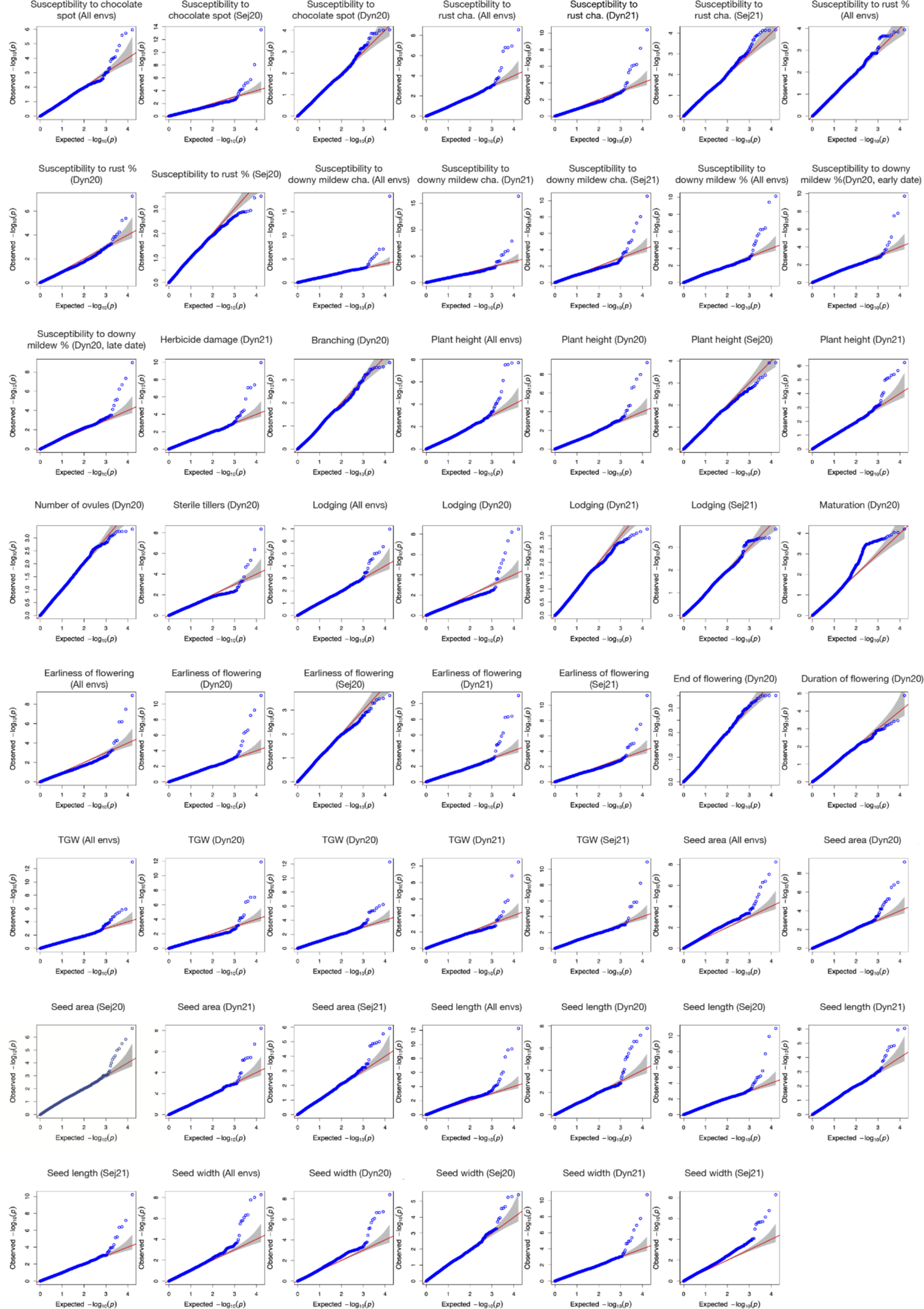
QQ-plots for all GWAS results. The plots show the observed distribution of *p*-values of markers tested for association in GWAS plotted against the expected distribution of *p*-values if no associated loci are found.

**Supplementary Figure 5.**
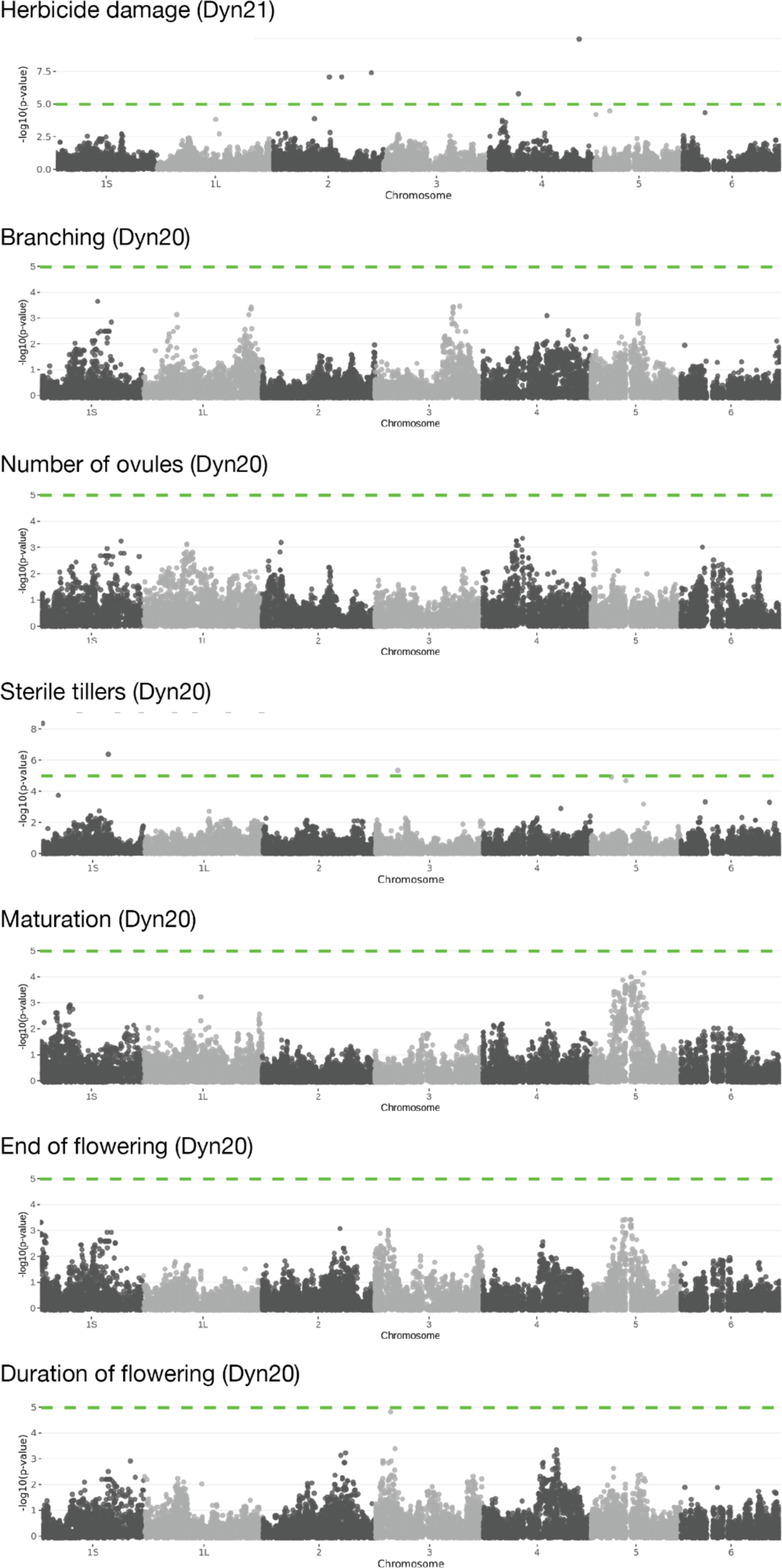
Additional Manhattan plots. Manhattan plots for GWAS of herbicide damage, branching, number of ovules, number of sterile tillers per plant, maturation date, end of flowering, and duration of flowering. The green line indicates the SimpleM-corrected threshold for significance.

**Supplementary Figure 6.**
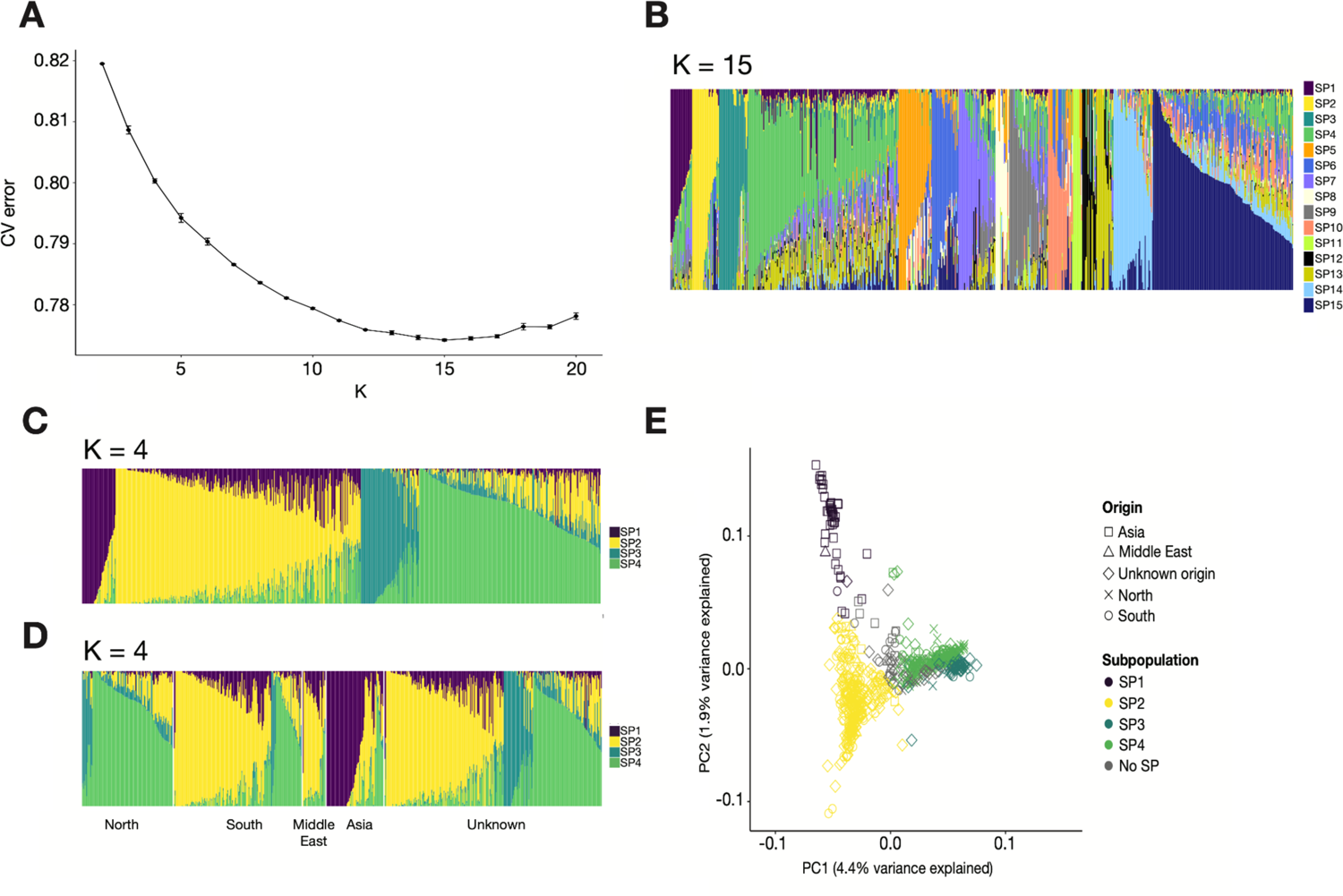
ADMIXTURE results. **A)** Cross-validation error of ADMIXTURE with K = 2 to K = 20. The bars display standard errors associated with repeating the CV 10 times for each value of K. B) ADMIXTURE proportions at K = 15 where the CV error is minimised. C–D) ADMIXTURE plots at K = 4. Each vertical bar represents a single accession coloured by its ancestry proportions. Accessions are grouped according to their subpopulation membership (C) or by their geographic origin (D). E) Principal component analysis (PCA) based on genotypes. The ADMIXTURE subpopulations at K = 4 are represented by colours and geographic origin is represented by shapes.

**Supplementary Figure 7.**
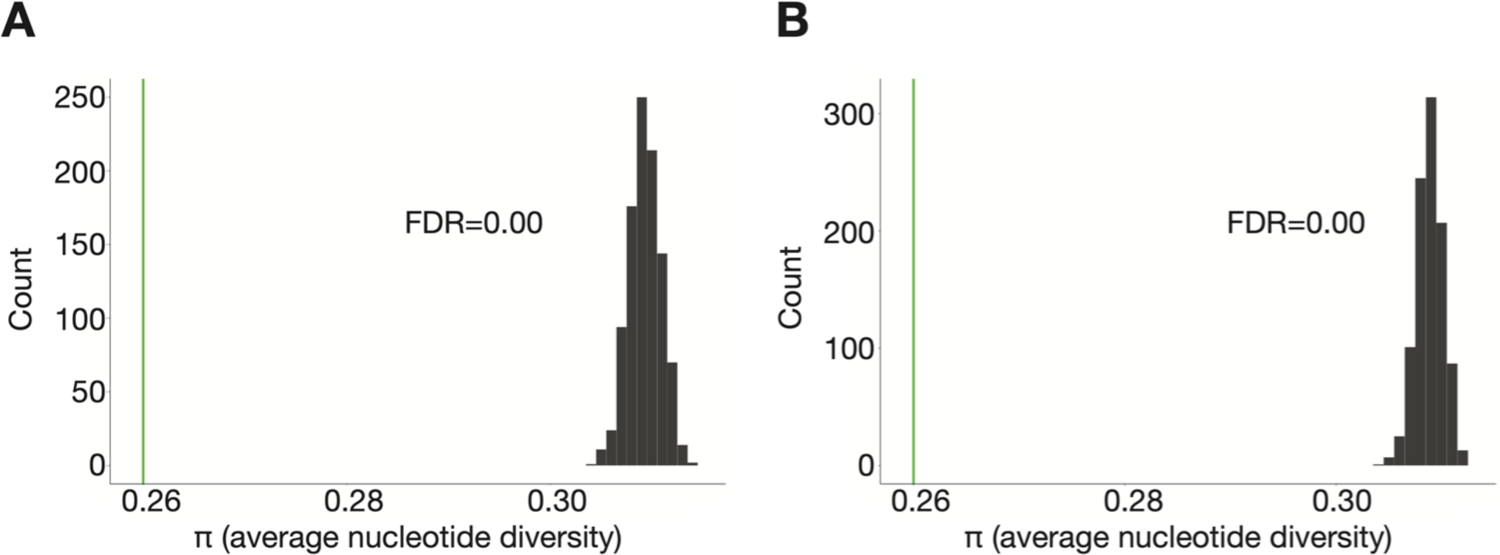
Distribution of average genome-wise nucleotide-diversity (π) values of 1000 subsets (n = 49) of SP1 (A) and SP2 (B). The horizontal green line displays the π value for SP3 (n = 49).

**Supplementary Figure 8.**
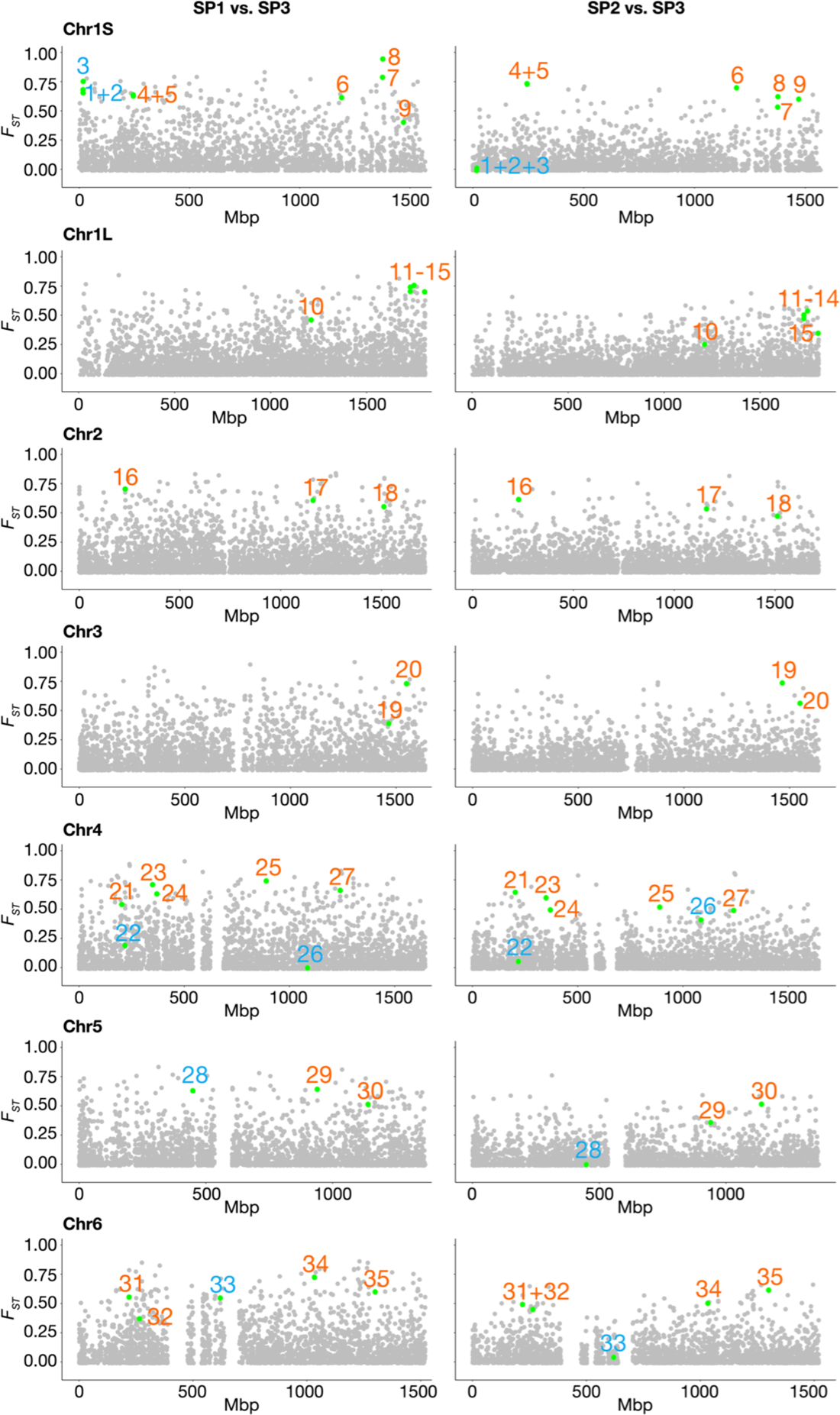
Genome-wide distribution of *F_ST_* values for pairs of subpopulations. The *F_ST_* values of each SNP throughout a chromosome are displayed as grey dots. The green dots report the 35 SNPs under selection identified in the outlier scans. The numbers next to the green dots serves as a marker code: 1: AX-416824401, 2: AX-416760427, 3: AX-416791399, 4: AX-416723873, 5: AX-416737096, 6: AX-416776470, 7: AX-181492359, 8: AX-416745027, 9: AX-416819371, 10: AX-416741889, 11: AX-181188041, 12: AX-181482613, 13: AX-416771656, 14: AX-416765862, 15: AX-181440418, 16: AX-181487950, 17: AX-181175939, 18: AX-181486832, 19: AX-181194098, 20: AX-416747475, 21: AX-416778737, 22: AX-416724016, 23: AX-416761735, 24: AX-181496895, 25: AX-416722420, 26: AX-181165197, 27: AX-416775196, 28: AX-416783057, 29: AX-416763147, 30: AX-416779502, 31: AX-416767699, 32: AX-181158030, 33: AX-181497981, 34: AX-416738786, 35: AX-181155942. Markers in LD group 1 are highlighted in blue.

**Supplementary Figure 9.**
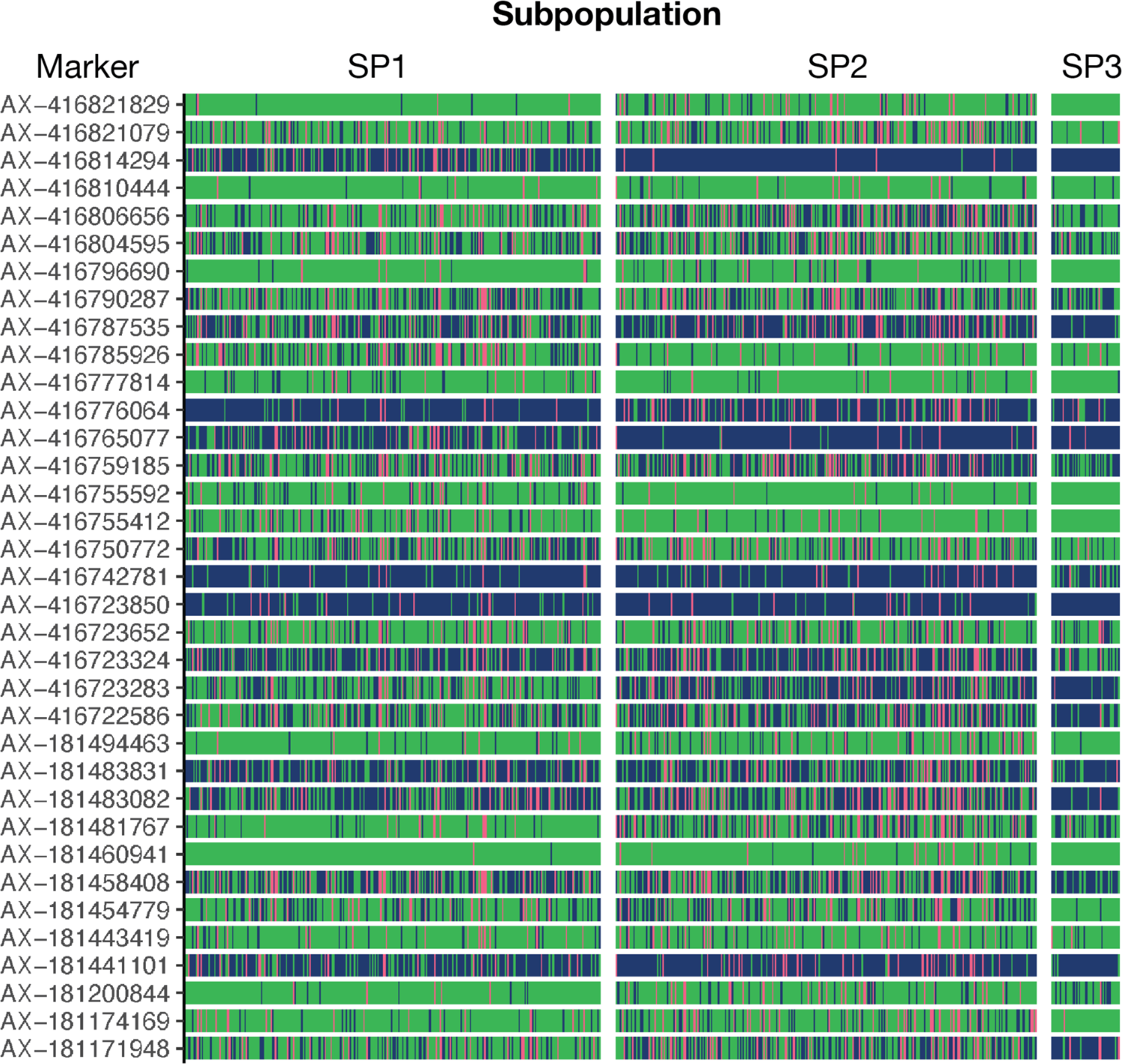
Segregation of 35 random markers. Each row shows the segregation pattern of one of 35 random markers. Each vertical line represents an accession and is coloured by genotype for a specific marker. Genotype coloring scheme is as follows: green, reference homozygote; pink, heterozygote; blue, alternative homozygote.

**Supplementary Figure 10.**
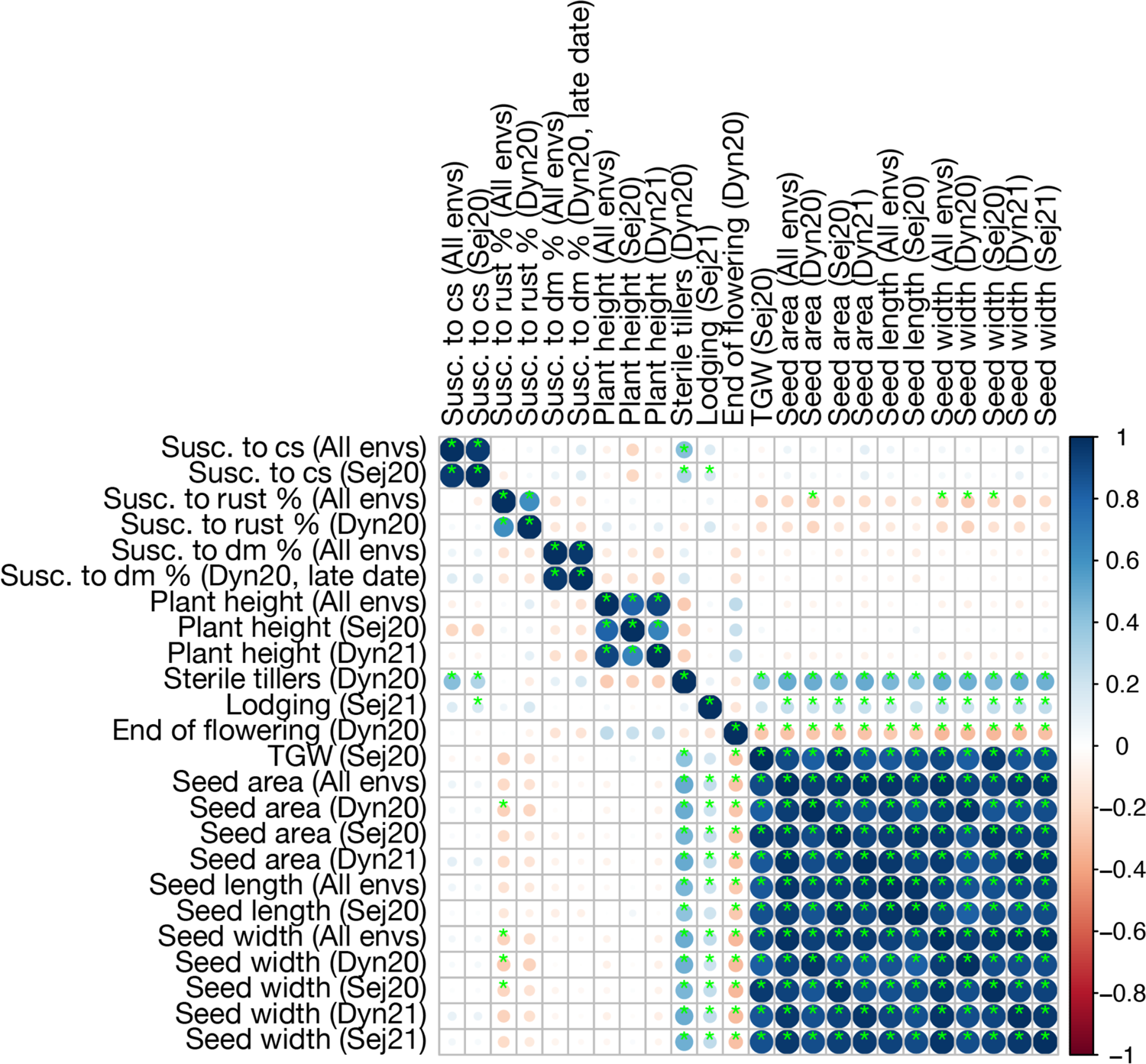
Genetic correlations between traits significantly explained by markers associated with North (SP1) versus South (SP2) differentiation. Green asterisk indicates statistical significance of correlation coefficients using a Bonferroni-corrected threshold of *p* < 0.05. Abbreviations: cha, character; cs, chocolate spot; dm, downy mildew; envs., environments; susc., susceptibility.

### Supplementary Tables

**Supplementary Table 1.**
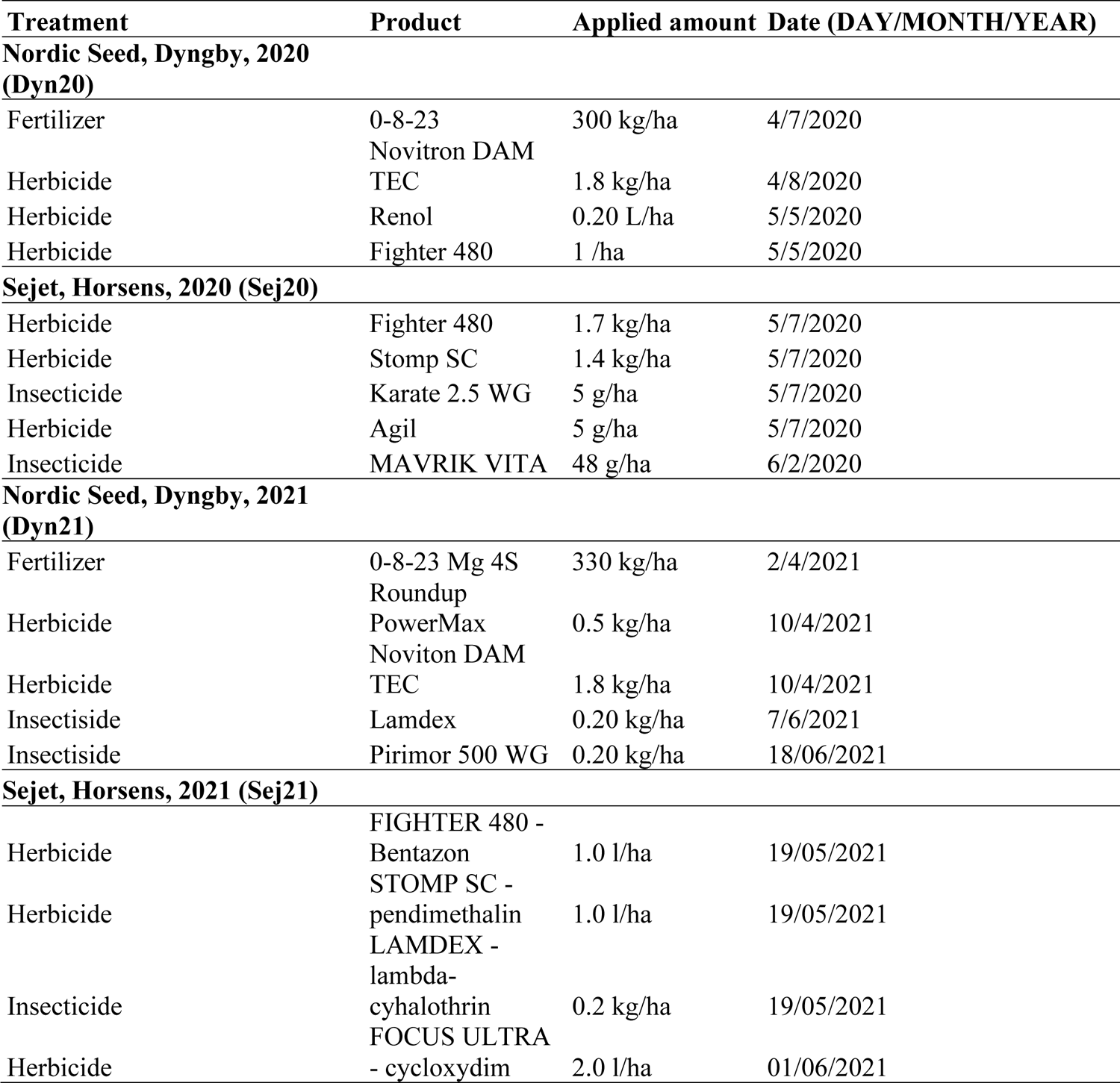
Management of trials.

### Supplementary files

**Supplementary File 1:** Passport information on individual accessions

**Supplementary File 2:** SNP markers and their positions

**Supplementary File 3:** Trait descriptions

**Supplementary File 4:** ANOVA results of the Seven-parent-MAGIC population

**Supplementary File 5:** Significant marker-trait associations identified by GWAS

**Supplementary File 6:** Individual results of all three methods of outlier detection

**Supplementary File 7:** Correlations between genetic values of all traits

**Supplementary File 8:** Seven-Parent MAGIC phenotypes used for GWAS

**Supplementary File 9:** Raw Phenotype Scores from Field Trials

